# A mutation-independent approach via transcriptional upregulation of a disease modifier gene rescues muscular dystrophy *in vivo*

**DOI:** 10.1101/286500

**Authors:** Dwi U. Kemaladewi, Prabhpreet S. Bassi, Steven Erwood, Dhekra Al-Basha, Kinga I. Gawlik, Kyle Lindsay, Ella Hyatt, Rebekah Kember, Kara M. Place, Ryan M. Marks, Madeleine Durbeej, Steven A. Prescott, Evgueni A. Ivakine, Ronald D. Cohn

**Affiliations:** Program in Genetics and Genome Biology, the Hospital for Sick Children Research Institute, Toronto, Canada; Department of Molecular Genetics, University of Toronto, Canada; Program in Neuroscience and Mental Health, the Hospital for Sick Children Research Institute, Toronto, Canada; Department of Physiology, University of Toronto, Canada; Unit of Muscle Biology, Department of Experimental Medical Science, Lund University, Lund, Sweden; Institute of Biomaterials and Biomedical Engineering, University of Toronto, Canada; Department of Pediatrics, the Hospital for Sick Children, Toronto, Canada

**Author notes:** These authors contributed equally to this work (Dwi U. Kemaladewi, Prabhpreet S. Bassi). These authors contributed equally to this work (Evgueni A. Ivakine, Ronald D. Cohn).

## Introductory paragraph

Identification of genetic modifiers has provided critically important insights into the pathogenesis and heterogeneity of disease phenotypes in individuals affected by neuromuscular disorders (NMDs). Targeting modifier genes to improve disease phenotypes could be especially beneficial in cases where the causative genes are large, structurally complex and the mutations are heterogeneous. Here, we report a mutation-independent strategy to upregulate expression of a compensatory disease-modifying gene in Congenital Muscular Dystrophy type 1A (MDC1A) using a CRISPR/dCas9-based transcriptional activation system.

MDC1A is caused by nonfunctional Laminin α2, which compromises muscle fibers stability and axon myelination in peripheral nerves ^1^. Transgenic overexpression of *Lama1*, encoding a structurally similar protein Laminin α1, ameliorates muscle wasting and paralysis in the MDC1A mouse models, demonstrating its important role as a disease modifier ^2^. Yet, upregulation of *Lama1* as a postnatal gene therapy is hampered by its large size, which exceeds the current genome packaging capacity of clinically relevant delivery vehicles such as adeno-associated viral vectors (AAVs).

In this study, we use the CRISPR/dCas9-based transcriptional activation system to upregulate *Lama1*. This system is comprised of catalytically inactive *S. aureus* Cas9 (dCas9) fused to VP64 transactivation domains and sgRNAs targeting the *Lama1* promoter. We first demonstrate robust upregulation of *Lama1* in myoblasts, and following AAV9-mediated intramuscular delivery, in skeletal muscles of *dy^2j^/dy^2j^* mouse model of MDC1A.

We therefore assessed whether systemic upregulation of *Lama1* would yield therapeutic benefits in *dy^2j^/dy^2j^* mice. When the intervention was started early in pre-symptomatic *dy^2j^/dy^2j^* mice, *Lama1* upregulation prevented muscle fibrosis and hindlimb paralysis. An important question for future therapeutic approaches for a variety of disorders relates to the therapeutic window and phenotypic reversibility. This is particularly true for muscular dystrophies as it has long been hypothesized that fibrotic changes in skeletal muscle represent an irreversible disease state that would impair any therapeutic intervention at advanced stages of the disease. Here, we demonstrate that dystrophic features and disease progression were significantly improved and reversed when the treatment was initiated in symptomatic 3-week old *dy^2j^/dy^2j^* mice with already-apparent hind limb paralysis and significant muscle fibrosis.

Collectively, our data demonstrate the feasibility and therapeutic benefit of CRISPR/dCas9-mediated modulation of a disease modifier gene, which opens up an entirely new and mutation-independent treatment approach for all MDC1A patients. Moreover, this treatment strategy provides evidence that muscle fibrosis can be reversible to some degree, thus extending the therapeutic window for this disorder. Our data provide a proof-of-concept strategy that can be applied to a variety of disease modifier genes and a powerful therapeutic approach for various inherited and acquired diseases.

Congenital muscular dystrophy 1A (MDC1A) is an autosomal recessive neuromuscular disease associated with a high degree of morbidity and mortality in early childhood ^1^. It is caused by mutations in the *LAMA2* gene encoding Laminin α2 chain (LAMA2), which interacts with the β1 and γ1 chains to form the heterotrimer Laminin-211, an extracellular matrix protein (reviewed in ^3^). Laminin-211 interacts with α-dystroglycan and α7β1 integrin in skeletal muscle and Schwann cells, providing the necessary roles such as survival and stability of myotubes, as well as proper neurite growth, axon myelination and migration of Schwann cells. Lack of Laminin-211 in MDC1A causes degeneration of skeletal muscle and impaired Schwann cell differentiation, resulting in a cascade of secondary events including apoptosis/necrosis of muscle fibers, inflammation, and fibrosis, which ultimately precipitate the disease. Despite significant advances in our understanding of MDC1A pathophysiology, currently, there is no cure.

Due to the genetic nature of the disease, gene therapy is a promising treatment option for MDC1A. The large size of *LAMA2* transcript, however, presents a challenge with respect to gene delivery. We have recently overcome this challenge by using CRISPR/Cas9 technology to correct a mutation in *Lama2* gene *in vivo* ^4^. We focused on *dy^2j^/dy^2j^* mouse model, which harbors a splice site mutation in *Lama2* causing spontaneous exon skipping and truncation of N-terminal domain of the protein ^5^. We developed an exon inclusion strategy to correct the splice mutation, leading to restoration of full-length Lama2. This study established the first direct correction of the primary genetic defect underlying MDC1A in an *in vivo* model.

To date, there are over 350 reported pathogenic nonsense-, missense-, splice site- and deletion mutations in the *LAMA2* gene. Given the variety of MDC1A-causing genomic alterations, CRISPR/Cas9-mediated gene editing strategies would require design and thorough analysis of multiple sgRNAs specific for each individual mutation. Moreover, safety concerns regarding CRISPR/Cas9’s potential mutagenic nature and the presence of off-target effects after gene editing remain, which together may prove to be challenging from a safety and regulatory point-of-view. In contrast, the attenuation of disease pathogenicity by targeted modulation of disease modifier gene expression would be a potentially safer alternative and beneficial to all individuals with MDC1A.

One of the strongest reported disease modifiers for MDC1A is Laminin-α1 protein, which is structurally similar to Laminin-α2 (**Fig. 1a**). However, Laminin-α1 is not expressed in skeletal muscles or Schwann cells. A series of studies previously demonstrated that transgenic expression of *Lama1*, encoding Laminin-α1, rescued both myopathy and peripheral neuropathy in *dy^2j^/dy^2j^* ^6^ and *dy^3K^/dy^3K^* ^2, 7-10^; the latter also showed increased lifespan. Although these studies firmly established compensatory function of Laminin-α1 in MDC1A, utilization of this modifier as a postnatal gene therapy is hampered by the size of *Lama1* cDNA, which exceeds the 4.7 kb packaging capacity of AAV vectors. Advances in CRISPR/Cas9 technologies have provided opportunities for regulating gene expression and creating epigenetic alterations without introducing DNA double-strand breaks (DSBs); commonly termed CRISPR transcriptional activation/inhibition system. The strategy utilizes nuclease-deficient Cas9 (dCas9), which is unable to cleave DNA due to mutations within the nuclease domains, but still retains the ability to specifically bind to DNA when guided by a single guide RNA (sgRNA) ^11,12^. Using the originally described *S. pyogenes* (*Sp*) dCas9 fused to multiple copies of the VP16 transcriptional activator, our group and others have demonstrated utilization of the CRISPR/dCas9 system to upregulate expression of modifier genes *in vitro* ^11-14^. A major challenge for *in vivo* applications lies in the large size of *Sp*dCas9 and its transcriptional activator fusion derivatives that exceed AAV genome packaging capacity. To accommodate this limitation, we sought to adapt the transcriptional upregulation system and utilize a significantly smaller Cas9 protein derived from *S. aureus* (*Sa*)^15^ to upregulate MDC1A modifier *Lama1*. We hypothesized that CRISPR/dCas9-mediated transcriptional upregulation of *Lama1* would be sufficient to compensate for the lack of *Lama2* and ameliorate disease phenotypes in *dy^2j^/dy^2j^* mice.

**Fig. 1.**
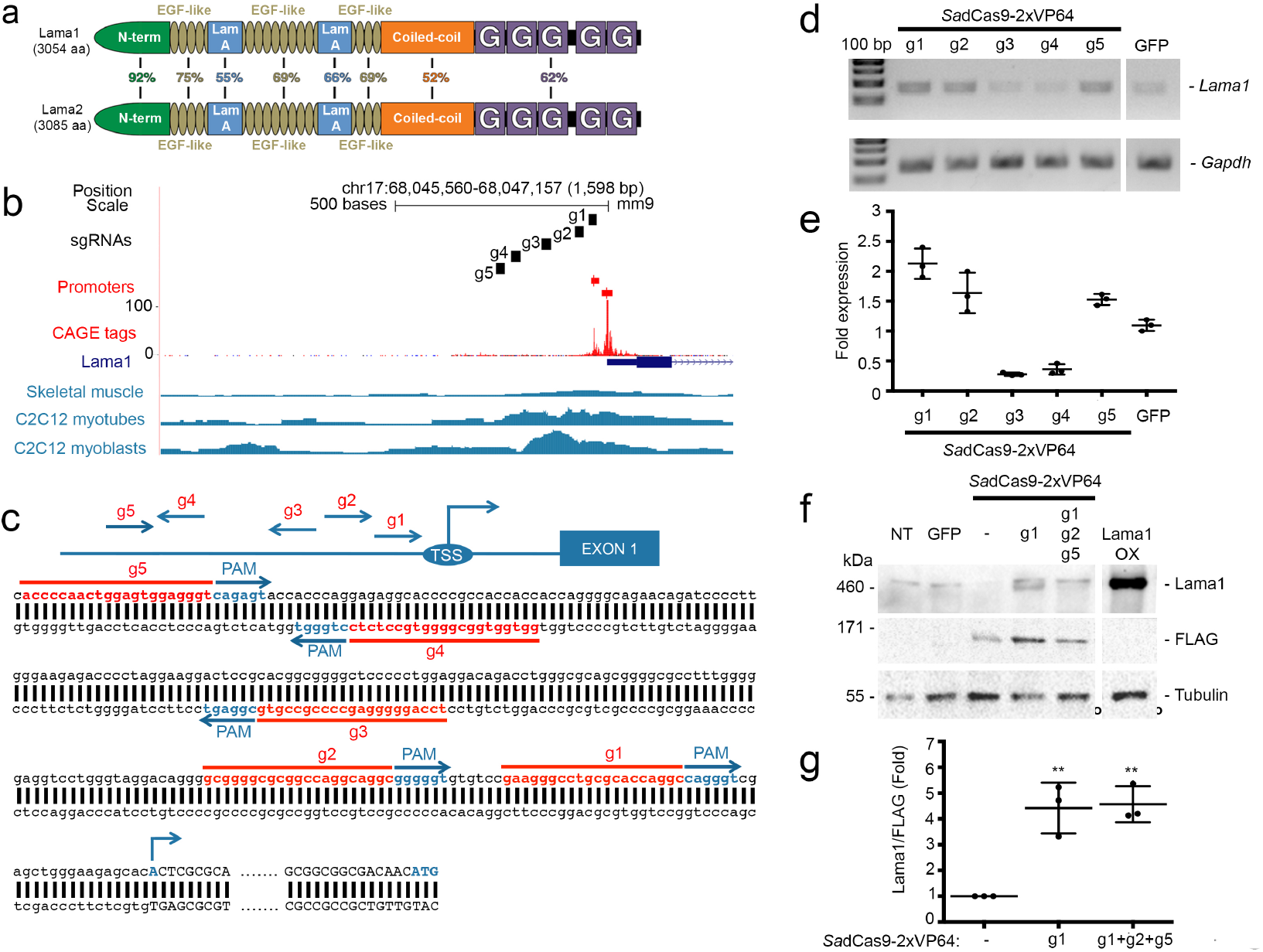
*Sa*dCas9-2xVP64-mediated upregulation of *Lama1 in vitro*. (**a**) Lama1 and Lama2 protein alignment. Total amino acid (aa) and percentage similarity between each domain are indicated. (**b**) Analyses of Lama1 proximal promoter. Five sgRNAs (g1-g5) were designed in the proximal promoter region of Lama1 immediately upstream of the transcription start site (TSS), as indicated by CAGE tags (red). Chromatin accessibility in skeletal muscle tissue and cells (retrieved using Digital DNase footprinting and ATAC-Seq) are shown in blue. Data were plotted according to positions from the UCSC Genome Browser. (**c**) Positions of the five sgRNAs relative to *Lama1* TSS. Arrowheads indicate the direction of each sgRNA. Sequences of each sgRNA (red) are immediately downstream (5’) of *Sa* PAM sequences (NNGRRT) (in blue). ATG indicates translation start site. (**d-e**) C2C12 myoblasts were transfected with a plasmid containing *Sa*dCas9-2xVP64 and the corresponding sgRNA(s) targeting *Lama1*, and 72 hours post-transfection, analyzed by (**d**) RT-PCR and (**e**) qRT-PCR. Single and combination of optimal sgRNAs were transfected into *dy^2j^/dy^2j^* myoblasts and Lama1 protein expression was assessed by western blot (**f**). FLAG expression serves as transfection control and used to (**g**) normalize Lama1 upregulation by densitometry analysis. Data are presented as mean ± standard deviation from at least three independent experiments. Statistical analysis was performed using Student’s *t-*test. \*\**P<*0.01.

First, we mutagenized *Sa*Cas9 endonuclease catalytic residues (D10A, N580A) to create *Sa*dCas9 and then fused it to transcriptional activators VP64 (four copies of VP16) on both N- and C-termini (**Fig. S1**). We tested the ability of the system, denoted as *Sa*dCas9-2xVP64, to upregulate the expression of minCMV-driven *tdTomato* gene in 293T cells ^16^. In the absence of the *Sa*dCas9-2xVP64, the expression of *tdTomato* was undetectable due to the low baseline activity of minCMV promoter (**Fig. S1a, b**). In the presence of *Sa*dCas9-2xVP64 in combination with an sgRNA targeting the minCMV locus, we observed high *tdTomato* fluorescence signal (**Fig. S1c, d**), indicating the general applicability of this system to modulate expression of a gene of interest. Subsequently, we tailored the system to upregulate *Lama1* expression and designed five sgRNAs, denoted as g1 to g5, within the 500 nucleotides region immediately upstream of the *Lama1* transcription start site (**Figs. 1b, c**). When co-expressed with *Sa*dCas9-VP64 in C2C12 murine myoblasts, 3 out of 5 sgRNAs, namely g1, g2, and g5 consistently induced significant increase of *Lama1* transcript expression (**Figs. 1d, e**).

Although all sgRNAs were designed to target a chromatin-accessible region derived from DNAse1 hypersensitivity- and assay for transposase-accessible chromatin (ATAC-Seq) sites (**Fig. 1b**), g3 and g4 failed to increase expression of *Lama1* above the basal level. This corroborates previously reported findings that chromatin accessibility is not a reliable predictor of successful gene activation ^12,17,18^. Nonetheless, the sgRNA closest to the transcription start site, g1, and the combination of the three most optimum sgRNAs (g1, g2, g5) resulted in a robust increase in Lama1 protein expression in *dy^2j^/dy^2j^*-derived myoblasts (**Figs. 1f, 1g**), warranting further investigation *in vivo*.

We then treated 3-week-old *dy^2j^/dy^2j^* mice with an AAV9 carrying FLAG-tagged *Sa*dCas9-2xVP64 in the absence of sgRNA (no guide) or with g1 (single guide) or a combination of g1, g2 and g5 (three guides) (**Fig. 2a**). Due to the packaging capacity of the AAV, the dCas9-2xVP64 and the three guides were split into two vectors (**Fig. 2a**). Each mouse received a single intramuscular injection of 7.5×10^11^ viral genomes of AAV9, which was doubled to 1.5×10^12^ viral genomes total for the three guide cohort, in the right *tibialis anterior* (TA) and was sacrificed 4 weeks post injection. FLAG expression was detected by western blot in all *Sa*dCas9-2xVP64-injected right TA muscles, however, only those injected with guide-containing constructs showed Lama1 upregulation (**Fig. 2b**). Similarly, immunofluorescence staining revealed sarcolemmal expression of Lama1 (**Fig. 2c**, upper), indicating proper protein localization. Furthermore, H&E staining exhibited considerably improved muscle architecture (**Fig. 2c**, lower) in the single- and three guide treatment groups, as compared to the no guide controls.

**Fig. 2.**
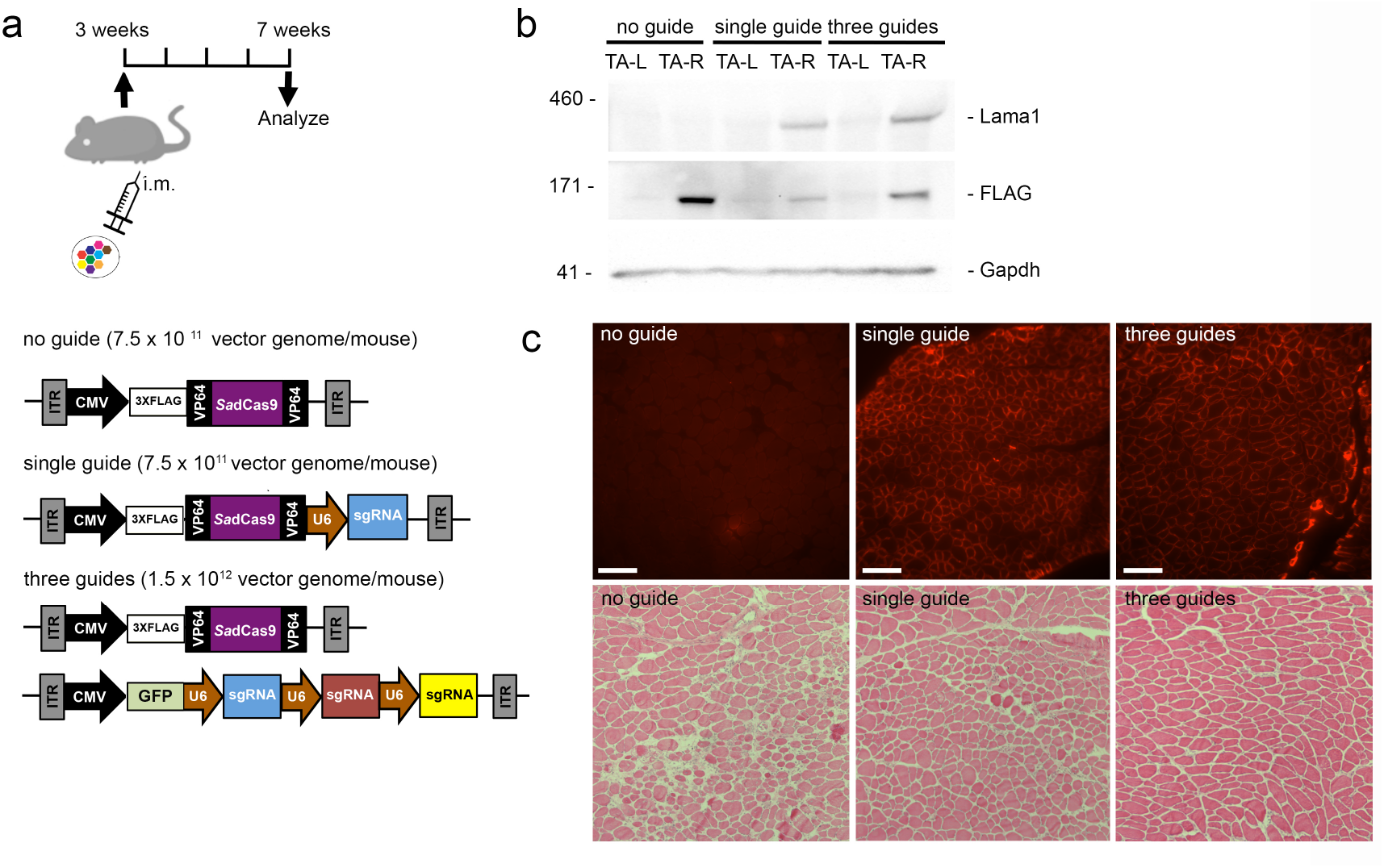
Upregulation of Lama1 in *tibialis anterior* (TA) muscles of *dy^2j^/dy^2j^* mice following intramuscular administration. (**a**) Right TA muscles of 3-week old *dy^2j^/dy^2j^* mice were injected with AAV9-carrying no guide (n=4; 7.5×10^11^ viral genomes), single guide (n=4; 7.5×10^11^ viral genomes) or three guides (n=4; split into two vectors, thus total dose was 2×7.5×10^11^ viral genomes). ITR: Inverted Terminal Repeats. CMV and U6 promoters are depicted in arrows. Left TA muscles serve as control. (**b**) Western blot analysis on Lama1, FLAG-tagged *Sa*dCas9, and Gapdh expression. (**c**) Immunofluorescence (upper) and H&E (lower) stainings on cross-sections of right TA muscles from each treatment group. Scale bar: 50 μm. Representative images from one animal per treatment group are shown.

Next, we sought to investigate whether upregulation of *Lama1* could be achieved systemically *in vivo* and, if administered into pre-symptomatic neonatal *dy^2j^/dy^2j^* mice, would prevent the manifestation of dystrophic pathophysiology and paralysis. AAV9 particles carrying either no guide or a combination of three guides were injected into the temporal vein of 2-day-old (P2) *dy^2j^/dy^2j^* mice (**Fig. 3a**). Seven weeks post injection, the animals that received the three guides treatment showed an absence of classical hindlimb contracture, unlike the control group (**Fig. 3b; Supplementary videos 1** and **2**). Western blot, immunofluorescence and H&E staining of TA and gastrocnemius muscles demonstrated robust expression of Lama1 on the sarcolemmal membrane (**Fig. 3c, 3d**), leading to ~50% reduction of fibrosis (**Figs. 3e, 3f**, **S2**).

**Fig. 3.**
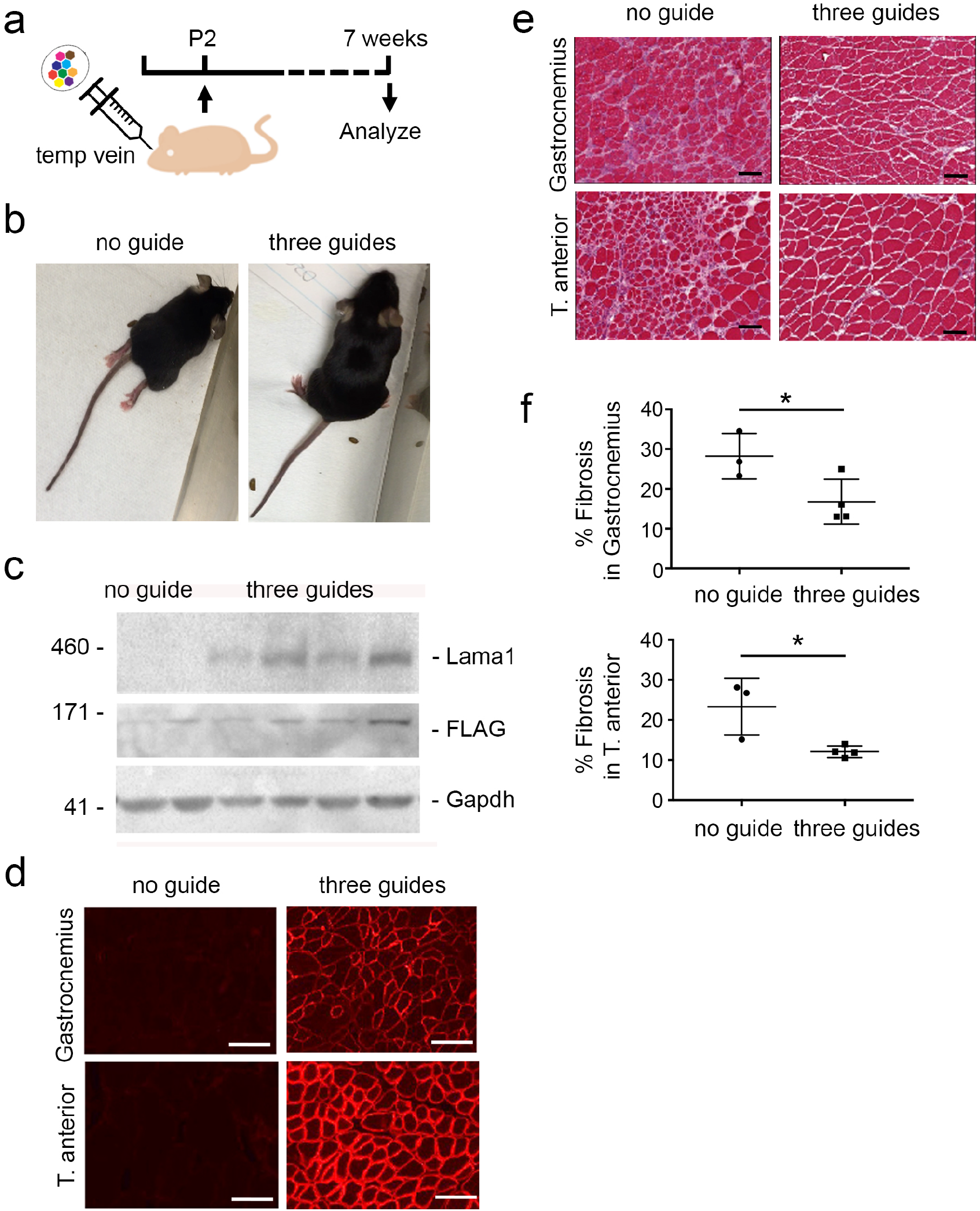
Early intervention to upregulate Lama1 prevents disease progression in *dy^2j^/dy^2j^* mice. (**a**) Two-day-old neonatal *dy^2j^/dy^2j^* mice were injected with AAV9 carrying no guide (n=3; 7.5×10^11^ viral genomes) or three guides (n=4; split into two vectors, thus total dose was 2×7.5×10^11^ viral genomes) via temporal vein and sacrificed 7 weeks later. (**b**) Photographs of *dy^2j^/dy^2j^* from both treatment groups prior to sacrificing. Note the difference in hindlimb contractures. Lama1 expression was analyzed by (**c**) western blot and (**d**) immunofluorescence staining, and general histopathology was evaluated by (**e**) H&E staining. The muscle groups analyzed are indicated on each panel, except on (**c**), which shows tibialis anterior (TA) muscles. Scale bars: 100 μm (**d**), 200 μm (**e**). (**f**) The percentage of fibrosis from gastrocnemius (upper) and TA (lower) muscles are calculated and presented as mean ± standard deviation. Statistical analysis was performed using Student’s *t-*test. \**P<*0.05.

We subsequently examined the ability of *Lama1* upregulation to reverse established muscular and peripheral nerve dysfunctions by initiating the treatment at 3 weeks of age, when paralysis was already apparent ^19,20^ (**Fig. 4a**). We first tested three different doses, ranging from 7.5×10^10^ to 3×10^11^ viral genome/gram of mouse per AAV9, which was doubled for the three guide cohorts due to the utilization of two vectors, and found that the highest dose resulted in homogeneous Lama1-positive fibers (**Fig. S3a**), significant improvement of muscle function and mobility (**Figs. S3b, S3c**). Longitudinal measurement of animal mobility and stand-up activity in a non-invasive open field activity assay showed significant difference between no guide and three guides-treated mice starting at 5 and 6 weeks old, respectively, which was sustained over time (**Figs. 4b, 4c**). Furthermore, specific tetanic force, which measures the aggregate torque produced by the dorsi flexor muscles, was also improved in the three guides-treated mice (**Fig. 4d**). In line with this finding, we observed a significant increase in nerve conduction velocity, which indicates restoration of the myelination defect and contributes to neuromuscular functionality (**Fig. 4e**). The absence of paralysis in the hind limbs and markedly improved movement of the mice were evident at the end of the treatment regimen (**Supplementary videos 3, 4**).

**Fig. 4.**
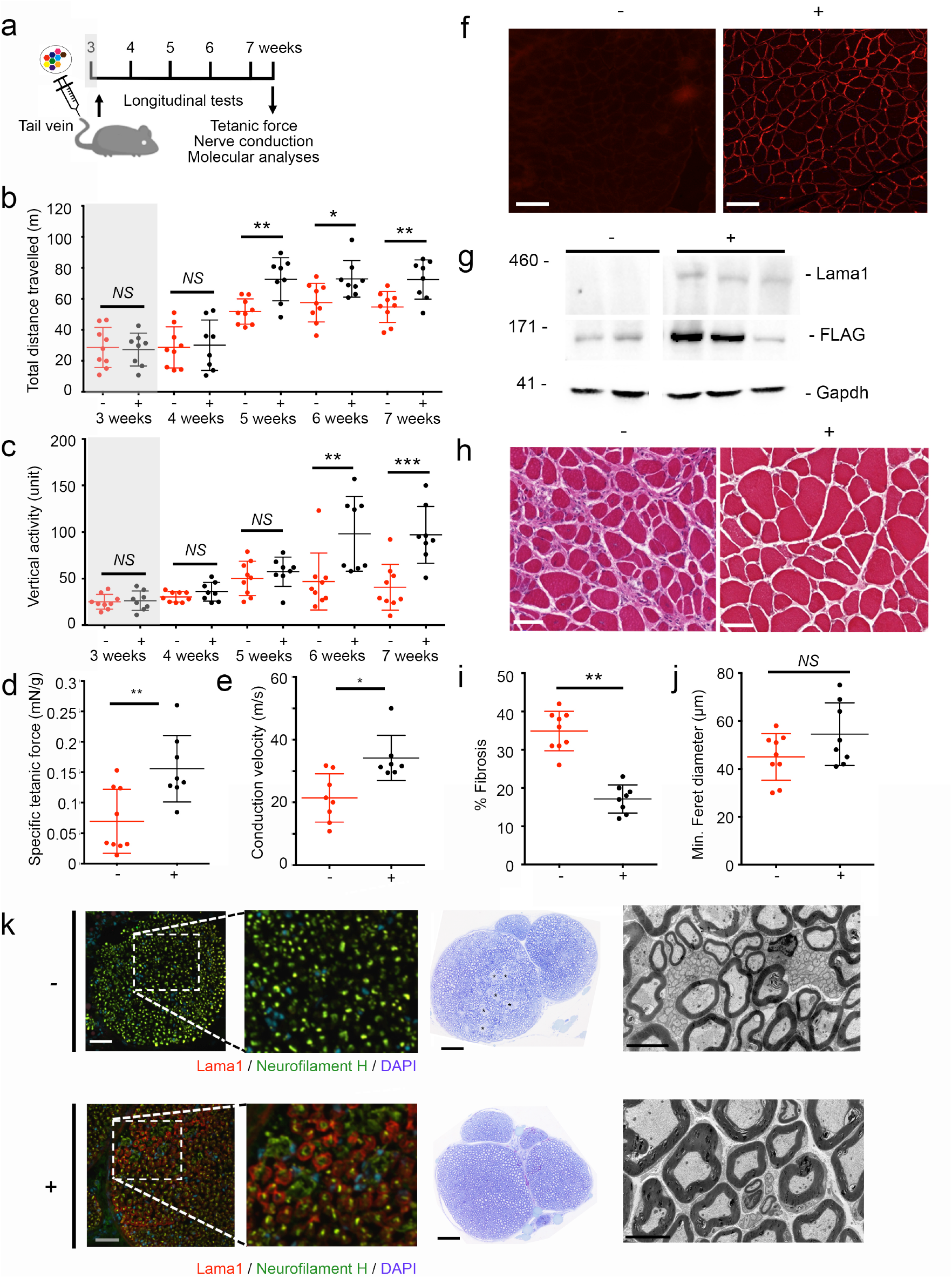
Upregulation of Lama1 in older *dy^2j^/dy^2j^* mice halts disease progression. (**a**) Three-week old *dy^2j^/dy^2j^* mice were injected with AAV9 carrying no guide (denoted as -; n=9; 3×10^11^ viral genomes/gram) or three guides (denoted as +; n=6; split into two vectors, thus total dose was 2×3×10^11^ viral genomes/gram) via tail vein. Grey box represents period before treatment. (**b, c**) Open field activity assay was performed weekly, before (grey-shaded) and after treatment, and the mice were tested for (**d**) muscle contractile and (**e**) nerve conduction velocity at the end of the treatment regimen, prior to molecular analyses of the tissues. (**f**) Immunofluorescence staining and (**g**) western blot to analyze Lama1 expression and (**h**) H&E staining to quantify (**i**) fibrosis and (**j**) fiber size were performed on gastrocnemius muscles. (**k**) Sciatic nerves were stained for Lama1 (red) and Neurofilament H (green) (left). Nuclei were counterstained by DAPI (blue). Higher magnification images from the dotted area are shown. Representative toluidine blue staining (middle) and electron micrograph (right) are shown. Asterisks indicate region of amyelinated axon fibers. Scale bars: 100 μm (**f**), 50 μm (**h**), and 200 μm (**k**). Data are presented as mean ± standard deviation. Statistical analysis was performed using Student’s *t-*test. \**P<*0.05. \*\**P<*0.01.

Molecular analysis of the treated mice revealed strong Lama1 expression by immunostaining and western blot (**Figs. 4f, 4g, S4**), which was accompanied by normalization of α4 chain of the laminin subunit (**Figs. S5, S6**) ^6^, significant improvement in muscle histopathology (**Fig. 4h**) and approximately 50% reduction in the fibrotic area (**Fig. 4i**) compared to the no guide control group. There was a trend towards larger fibers in the treated mice, although it did not reach statistical significance due to the large variation between animals (**Fig. 4j**). In addition, upregulation of Lama1 was also observed in the endoneurium of sciatic nerves and resulted in restoration of myelination defect (**Figs. 4k, S7**), supporting the improvement of nerve conduction velocity and lack of paralysis in the mice (**Fig. 4e**, **Supplementary videos 3, 4**). Quantification of the AAV genome copy number revealed accumulation of most of the viral genome in the liver, which is expected from intravenous delivery. Nevertheless, approximately 1.11±0.4 and 34.1±4.3 copies/diploid genome of the viral genome were detected in the sciatic nerves and skeletal muscles, respectively (**Fig. S8**). Remarkably, even relatively low transduction efficiency in sciatic nerves was sufficient for functionally significant upregulation of Lama1 expression.

Finally, we investigated the genome wide effects of CRISPR/dCas9-mediated Lama1 upregulation by performing RNA-sequencing on quadriceps muscles isolated from mice treated with AAV9 carrying no guide and three guides (**Figs. S9, S10, Supplemental Tables 1-3**). Age-matched wildtype and *dy^2j^/dy^2j^* mice served as controls. We observed a 3.6-log_2_fold upregulation of Lama1 (defined by a false discovery rate, FDR <0.05) when comparing the mice treated with AAV9 carrying no guide and three guides (**Fig. S9a**). The transcriptional change was even higher at 9-log_2_fold when comparing untreated *dy^2j^/dy^2j^* with three guide-treated *dy^2j^/dy^2j^* (**Fig. S9b, S9c**). Hierarchical clustering between groups revealed clustering between wildtype and three guides, whereas the untreated cohort was clustered together with no guide-treated mice (**Figs. S9d, S10**). We also computationally predicted 704 potential off-target binding sites for the three sgRNAs targeting *Lama1* promoter in the mouse genome, selected based on the presence of a 6bp PAM-proximal seed sequence and fewer than ten total mismatches to the cognate target sequence (**Supplemental Tables 4-5**). None of the top 100 genes contained a predicted off-target site within 60 kilobases of the gene body, with the average distance from off-target site to the gene body being 2.2 megabases. Taken together, our results provide strong evidence of the robustness and durability of CRISPR/dCas9-mediated Lama1 upregulation for the treatment of MDC1A. Additionally, we show the therapeutic promise of the strategy for reversal of dystrophic feature in skeletal muscle and peripheral neuropathy in *dy^2j^/dy^2j^* mouse model of MDC1A, which ultimately halts progression of the disease, without any confounding off-target effects.

One challenge in developing a therapy for MDC1A is that the heterogeneity of mutations often leads to variable disease severity and progression. Therefore, there is an urgent need to develop a universal, mutation-independent strategy that provides a treatment approach for all patients with MDC1A. Our study establishes a framework in which CRISPR/dCas9 transcriptional upregulation of a disease modifier gene, such as *Lama1,* ameliorates disease symptoms *in vivo* and has the potential to be applied to all MDC1A patients, irrespective of their mutations.

Advances have been made towards elucidating MDC1A pathogenesis due to the availability of several mouse models with absent or reduced *Lama2* expression. Yet, an important question in the development of therapeutics and clinical trials in MDC1A is the reversibility of symptoms caused by muscle fibrosis and nerve abnormalities. This issue has been challenging to address in patients^21^, however, the ability to modify gene expression in postnatal animals allowed us to address this question and begin to investigate the therapeutic window of intervention.

We have previously demonstrated that an early intervention using CRISPR/Cas9-mediated correction of a splicing defect resulted in robust Lama2 restoration and prevention of disease manifestation ^4^. Here we showed that upregulation of *Lama1*, when initiated at pre-disease-onset, leads to similar prevention. Importantly, when the therapeutic intervention was initiated at older age, significant rescue of the phenotypes was attainable, indicating that post-symptomatic treatment provides a significant benefit in the *dy^2j^/dy^2j^* mouse model.

In addition to *Lama1* upregulation described in this study, a number of disease modifying strategies are currently being explored in MDC1A animal models, including treatment with miniaturized agrin ^22-24^ and laminin-α1 LN-domain nidogen-1 (αLNNd) ^25-27^. While the efficacy of αLNNd has only been explored in transgenic mice, AAV-mediated delivery of mini agrin has been shown to normalize most histopathological parameters in skeletal muscle, and improve myelination and regeneration of Schwann cells of the peripheral nerves ^22,24^. Despite the observed phenotypic improvement, the mini agrin-treated mice still have a lower survival rate compared to wild-type animals expressing full-length agrin, suggesting the potential shortcoming of the shortened version of agrin, which may be overcome by its full-length form in native glycosylation state ^25^. It is important to note that many preclinical studies in MDC1A were carried out in the more severe *dy^W^/dy^W^* mouse model. Therefore, as a follow up on our study, it will be important to evaluate our approach in the *dy^W^/dy^W^* mice to assess critical parameter such as survival, which is not possible in the *dy^2j^/dy^2j^* mouse model due to its near-normal life span. Overall, our approach in combining the CRISPR/dCas9-mediated transcriptional upregulation and AAV9 as a delivery vehicle can be translated to many other disease modifiers, such as agrin, as well as conditions where modulation of disease modifiers is required within both skeletal muscles and peripheral nerves.

In fact, neuromuscular disorders have provided excellent examples to demonstrate the role of disease modifiers (recently reviewed in ^28^). Beyond MDC1A, several studies have demonstrated that upregulation of *Lama1* stabilizes the sarcolemmal membrane in dystrophin-deficient mouse models ^29^. The most advanced approach is via delivery of Laminin-111 protein, although the efficacy remains low and the need to produce a large amount of bioactive protein is challenging ^29^. In addition, the utilization of CRISPR/dCas9 system to upregulate Lama1 in *mdx* mice has been achieved locally via electroporation, which is not easily translatable into clinical settings ^30^. Our strategy of employing AAV-mediated *S. aureus* dCas9 to upregulate *Lama1 in vivo* may be tested further as a potential therapy in the context of Duchenne muscular dystrophy.

In addition, since the CRISPR/dCas9-mediated transcriptional modulation acts directly on the endogenous locus, it may be used in conjunction with a mutation-correction approach where the level of restoration of the defective gene is suboptimal, therefore necessitating further amplification to reach therapeutic efficacy ^4, 31-36^.

A very recent study by Liao *et al* described utilization of the CRISPR/Cas9 system to recruit MCP:P65:HSF1 transcriptional activation complex to induce expression of target genes in skeletal muscle, kidney and liver tissues ^18^. This resulted in phenotypic augmentation such as enhanced muscle mass and substantial improvement in disease pathophysiology, thereby highlighting the feasibility of using CRISPR/dCas9-mediated transcriptional activation as a possible therapeutic modality. However, their study relied almost exclusively on a Cas9-expressing transgenic mouse model or local intramuscular treatments, and therefore it is difficult to extrapolate the efficacy of this strategy to disease-relevant models. In contrast, we successfully demonstrated robust upregulation of Lama1 after systemic delivery of therapeutic components in a relevant mouse model of disease that does not constitutively express Cas9.

Finally, the modular nature of the CRISPR/dCas9 system can be utilized to not only to upregulate, but also to downregulate target gene expression. The latter can be achieved by coupling dCas9 with transcriptional repressor such as Kruppel-associated box (KRAB) ^17,37^. A very recent study described that following sarcolemmal injury, the muscle membrane resealing process is greatly improved upon the deletion of Osteopontin, which acts in a concerted fashion with protective modifiers such as Latent TGF-β binding protein (LTBP4) and Annexins 1 and 6 ^38,39^. The combinatorial effects of such modifiers, whether they are additive, synergistic or even opposing in action, represent a new paradigm for lessening disease phenotypes. A foreseeable application of CRISPR/dCas9-mediated modulation is in the upregulation of protective disease modifier genes, such as Lama1 or LTBP4, with concurrent downregulation of detrimental genes, such as Osteopontin, providing a combinatorial therapeutic approach.

In summary, our study establishes a framework to utilize CRISPR/dCas9 to modulate gene expression of disease modifiers that should be considered as a mutation-independent therapeutic strategy not only to MDC1A, but also to various other inherited and acquired diseases.

## Methods

### Engineering of activation constructs

A fragment containing a catalytically inactive *SaCas9* coupled to two flanking VP64 transactivator domains was synthesized by BioBasic Canada and cloned into pX601 (Addgene 61591) using AgeI and EcoRI directional cloning to generate 3XFLAG-VP64- SadCas9 (D10A/N580A)-NLS-VP64 plasmid (**Figs. 2a, S11, Supplemental Table S6**). Each sgRNA **(Supplemental Table S6)** was subsequently introduced using BsaI directional cloning. To generate the three guides only construct (**Fig. 2a**), a fragment containing three repetitive regions of U6 promoter and *S. aureus* guide scaffold was assembled, with short linkers in between each region (BioBasic Canada). The fragment was cloned into KpnI and NotI sites of a pX601-derivative plasmid.

### Cell Culture

Primary myoblasts were isolated from the Extensor Digitorum Longus (EDL) Muscle of *dy^2J^/dy^2J^* mice, as previously described ^40^ and maintained in DMEM supplemented with 1% chicken embryo extract (GeminiBioscience), 10% horse serum, 1% penicillin/streptomycin and 1% L-glutamine (all from Gibco, unless indicated otherwise). HEK293 and C2C12 cells where maintained in DMEM supplemented with 10% FBS, 1% penicillin/streptomycin and 1% L-glutamine (all from Gibco). All cells were maintained at 37°C with 5% CO_2_.

Transfection of HEK293T cells was performed as previously described^2^. C2C12 and *dy^2J^/dy^2J^* cells were transfected in 12-well plates using the Neon Transfection System (Invitrogen). 400,000 cells were electroporated with 1.5 μg of DNA utilizing optimization program 16 (pulse voltage: 1400V, pulse width: 20ms, pulse number: 2). Cells were grown for 72 hours, after which RNA or protein was subsequently collected for protein analysis and guide screening.

### Animals, virus production and injections

*dy^2j^/dy^2j^* mice were purchased from the Jackson Laboratory and maintained in the Toronto Center for Phenogenomics. Both male and female were used in the analyses. All animal experiments were performed according to Animal Use Protocol number 20- 0305. *Sa*dCas9-2xVP64, single guide and three guides plasmids (**Fig. 2a, Supplemental Table S6**) were packaged into AAV9 vectors by Vigene Biosciences as previously described ^3^. For intramuscular and temporal vein injection into neonatal pups, the dose of 7.5×10^11^ viral genomes each was used. Due to the limitation in packaging capacity, two AAVs were needed for the three guides cohort (**Fig. 2a**), therefore the total virus injected was 1.5×10^12^ viral genomes per animal. Injection volume was brought to 50 μl with 1XPBS (Gibco).

For the tail vein injection in young, 3 week old mice, three different doses were initially tested: 7.5×10^10^, 1.5×10^11^ and 3×10^11^ viral genomes per gram of mouse (**Fig. S3**). Similar to the intramuscular and temporal vein injections, two AAVs were needed for the three guides cohort, therefore the total dose used in experiments described in **Figs. 4, S4-S10** was 6×10^11^ viral genomes per gram of mouse. Injection volume was brought to 100 μl with 1XPBS (Gibco).

### RNA isolation, guide screening and RT-PCR

RNA was isolated from cultured cells and mouse tissue sections, and cDNA synthesis was performed as previously described^3^. PCR amplification was utilized to assess the efficiency of each guide in upregulating *Lama1* expression using a primer in *Lama1* exon 55 (RDC 1919) and a second primer spanning the junction of exons 55 and 56 (RDC 1920). Sequences are listed in **Supplemental Table S6**.

qPCR utilizing Fast SYBR green Master Mix (Qiagen) on a Step One Plus Real Time PCR (Applied Biosystems) was performed. *Lama1* expression was analyzed using a primer in *Lama1* exon 55 (RDC 1919) and one spanning the junction of exons 55 and 56 (RDC 1920). Primers against endogenous Gapdh (RDC 345 and 346) were used as an internal control. ΔΔCt was analyzed to assess fold changes between treated and untreated samples.

### Protein Isolation and western blot

Protein was isolated from *dy^2j^/dy^2j^* myoblasts and C2C12 cells by adding 150 μl of a 1:1 part solution of RIPA homogenizing buffer (50-mM Tris HCl pH 7.4, 150-nM NaCl, 1-mM EDTA) and RIPA double-detergent buffer (2% deoxycholate, 2% NP40, 2% Triton X-100 in RIPA homogenizing buffer) supplemented with protease-inhibitor cocktail (Roche).

Cells were subsequently scraped from the bottom of each well, collected and incubated on ice for 30 min. Cells then centrifuged at 12000xg for 15 min at 4°C and the supernatant was collected and stored at −80°C. Protein from mouse tissue sections was collected as previously described^3^. Whole protein concentration was measured using Pierce BCA protein assay kit according to the manufacturer’s protocol (Thermo Fisher Scientific). Western blot was performed as previously described^3^. Primary antibodies used were rabbit Anti-LNα1 E3 (a gift from Dr. Peter Yurchenco, 0.6 μg/ml), mouse monoclonal M2 anti-Flag (Sigma Aldrich F1804, 1:1000) and rabbit polyclonal anti-GAPDH (Santa Cruz sc-25778, 1:5000).

### Immunofluorescence and H&E staining

Muscles and nerves were sectioned at 8 μm thickness and processed for immunofluorescence analyses according to standard procedures. Antibodies used for immunofluorescence staining were rat monoclonal against Laminin α1 (mAb200, Durbeej Lab, 1:20), α2 (4H8-2, Sigma, 1:500), γ1 chain (clone A5, Thermo Fisher Scientific), rabbit polyclonal against Laminin α4 chain (kindly provided by Dr. Sasaki), mouse monoclonal against NF-H (Biolegend SMI 31, 1:1000), goat polyclonal anti-rat Alexa Fluor 555 (Thermo Fisher Scientific, 1:250) and goat polyclonal anti-mouse Alexa Fluor 488 (Thermo Fisher Scientific, 1:250). H&E staining was performed as previously described ^4^. Both immunofluorescence and H&E slides were scanned with the 3Dhistech Pannoramic 250 Flash II digital scanner and analyzed with CaseViewer software, with the exception of **Figs. S5**, **S6,** which were analyzed with a Zeiss Axioplan fluorescence microscope (Carl Zeiss) and images were captured using an ORCA 1394 ER digital camera (Hamamatsu Photonics) and Open Lab software version 4 (Improvision).

### Toluidine blue staining and electron microscopy

Freshly isolated mouse sciatic nerves were halved and fixed in a solution of 2% paraformaldehyde and 2.5% glutaraldehyde in 0.1M sodium cacodylate buffer until further use. For embedding, the specimens were rinsed with the 0.1M sodium cacodylate buffer, post-fixed in 1% osmium tetroxide in the washing buffer, dehydrated in a graded ethanol series followed by propylene oxide, and embedded in Quetol-Spurr resin overnight at 65^0^C. Sections of 90nm thickness were cut on a Leica EM UC7 ultramicrotome, stained with uranyl acetate and lead citrate, and imaged on a FEI Tecnai 20 electron microscope at 4400-, 10,000-, and 44,000X magnifications. The same 90nm sections were stained with toluidine blue and imaged on a Leica DM-2000. All reagents were purchased from Electron Microscopy Sciences. Quantification of myelin thickness was measured using ImageJ from at least 14 random axons per field (at least 5 fields per animal) ^41^.

### Open field and *in vivo* muscle force assays

Open field activity test and assessment of *in vivo* muscle force were performed on tail vein injected cohorts at the Lunenfeld-Tanenbaum Research Institute’s Centre for Modeling Human Disease Mouse Phenotyping Facility. For the open field test, mice were placed in the frontal center of a transparent Plexiglas open field (41.25 cm × 41.25 cm × 31.25 cm) illuminated by 200 lx. A trained operator, who is unaware of the nature of the projects and treatments, performed experiments. The VersaMax Animal Activity Monitoring System recorded vertical activities and total distance travelled for 20 minutes per animal.

*In vivo* muscle contraction test was performed using 1300A: 3-in-1 Whole Animal System and analyzed using Dynamic muscle control/analysis (DMC/DMA) High throughput software suite (Aurora Scientific). The mice were anaesthetised with intraperitoneal injection of ketamine/xylazine cocktail at 100 mg/kg and 10 mg/kg of body weight, respectively. Contractile output was measured via percutaneous electrodes that stimulate specific nerves innervating the plantar flexors. Specific tetanic force (200 ms of 0.5-ms pulses at 125 Hz) was recorded and corrected to body weight.

### Nerve conduction velocity

Mice were anesthetized with urethane (1.2 mg/g i.p.) and underwent a tracheotomy to maintain their airway but were not artificially ventilated. Core body temperature was maintained at 34-36.5°C using a feedback-controlled heating pad (TR-200; Fine Science Tools). The sciatic nerve was exposed at two locations via incisions (~15 mm) above the left knee (site #1) and along the sacral region of the vertebral column (site #2). After separating the nerve from adjacent tissue, a bipolar hook electrode was applied to the nerve at each site. Each electrode comprised two chlorided silver wires (0.01″ diameter, A-M systems) placed ~1 mm apart and bent at the tip to form hooks, and attached to a stimulator (Model DS3, Digitimer Ltd). Stimulating electrodes were insulated from underlying tissue using a small piece of plastic paraffin film. Stimulus duration was fixed at 20 μs and current intensity was varied. Stimulus timing was controlled by computer using a Power1401 computer interface and Signal v5 software (Cambridge Electronic Design, CED). The compound muscle action potential (CMAP) was recorded using needle electrodes (~30 G, BD *PrecisionGlide*™), one inserted into the gastrocnemius muscle and the other (reference) electrode inserted into the Achilles tendon. The CMAP signal was amplified, low-pass filtered at 10 Hz and high-pass filtered at 1 kHz (DAM 80, World Precision Instruments), and digitized at 40 kHz using the Power1401 and Signal v5 software (CED). To calculate conduction velocity (CV) from CMAP responses, the sciatic nerve was stimulated at just-maximal intensity (beyond which there is no change in CMAP amplitude or latency) three times at site #1 and again three times at site #2. Nerve conduction velocity was calculated as the difference between average CMAP latencies for each stimulation site divided by the length of nerve (7.5-10 mm) separating the sites.

### Vector genome quantification

Evaluation of AAV genome distribution was performed as previously published ^42^ with a few modifications. Genomic DNA was extracted from tibialis anterior, sciatic nerve, and liver of treated mice using Qiagen Blood and Tissue Kit. 90 ng of DNA was amplified using primers located in between the two inverted tandem repeats (ITRs) (RDC 1687, RDC 1679, **Supplemental Table S7**) using Fast SYBR green master mix (Qiagen) on a Step One Plus Real Time PCR (Applied Biosystems). The Ct value of each reaction was converted to viral genome copy number by interpolating against the copy number of standard curve of a known plasmid containing the sgRNA cassette (RDC 362). The amount of DNA input (90 ng per tissue) was used as a conversion factor to diploid genome (7 picogram DNA = 1 diploid genome).

### RNA-Sequencing

Total RNA was isolated from quadriceps muscles using RNeasy kit (Qiagen) and quantified using Qubit RNA HS assay (Thermo Fisher Scientific). RNA-Sequencing was performed by the Centre for Applied Genetics in Toronto using the Illumina HiSeq 2500 system, producing 120bp paired-end reads. Raw transcript reads were aligned to the GRCm38 mouse genome (mm10) using HISAT2 ^43^. HTSeq was used to determine the absolute number of read counts for each gene ^44^. Only genes with at least 1 read per million in at least three replicates were kept for downstream analysis. Normalization and differential expression analysis was completed using the R packages limma, v3.36.3 and edgeR, v.3.22.3 ^45^. Differentially expressed genes were defined as genes with a more than twofold change and adjusted *P* < 0.05. Off-target analysis was conducted using a list of 704 computationally predicted using Cas-OFFinder ^46^. Using the Bedtools suite “closest” function, each of the top 100 differentially expressed genes was matched to the nearest off-target loci to determine the shortest distance from off-target loci to differentially expressed gene body.

### Statistical Analysis

GraphPad Prism (GraphPad software) was utilized to preform all statistical analyses. Two-tailed Student’ t-tests to evaluate statistical significance between two groups was preformed. Significance was considered to be P < 0.05.

### Data availability

The authors declare that the main data supporting the findings of this study are available within the article and its Supplementary Information files. Extra data are available from the corresponding authors upon request.

## Acknowledgements

The Cohn lab members are gratefully acknowledged for the technical support and critical input in this study. We thank P. Yurchenco (Rutgers University) for providing critical reagents in this study, C. Rand (Aurora Scientific), I. Vukobradovic (Clinical Phenotyping Core, Centre for Modeling Human Disease), R. Smith (Prescott Lab) for assistance with functional and behavioral studies, D. Holmyard (Nanoscale Biomedical Imaging Facility) for electron microscopy analysis and S. Pereira (The Centre for Applied Genomics) for genomic data acquisition.

This work was supported by AFM-Telethon, Cure CMD, Muscular Dystrophy Association (to D.U.K), Rare Disease Foundation Microgrant (to P.S.B. and S.E.), SickKids Restracomp (to S.E.), CIHR Summer Studentship (to R.M.), Natural Sciences and Engineering Research Council of Canada, Canadian Institute for Health Research, SickKids Foundation and R.S. McLaughlin Foundation Chair in Pediatrics (to R.D.C).

## Author contributions

D.U.K., E.A.I., R.D.C. conceived the study, D.U.K., P.S.B., S.E., D.A.B., K.I.G., K.L., E.H., R.K., K.M.P., R.M., performed the experiments, D.A.B., K.I.G., M.D., S.A.P. provided critical reagents, D.U.K., P.S.B, S.E., D.A.B, K.I.G, S.A.P., E.A.I, R.D.C. analyzed data, D.U.K, E.A.I., R.D.C. supervised the study, D.U.K. wrote the manuscript with inputs from the other authors. All authors provided feedback and agreed on the final manuscript.

## Extended data/supplementary figures

**Fig. S1.**
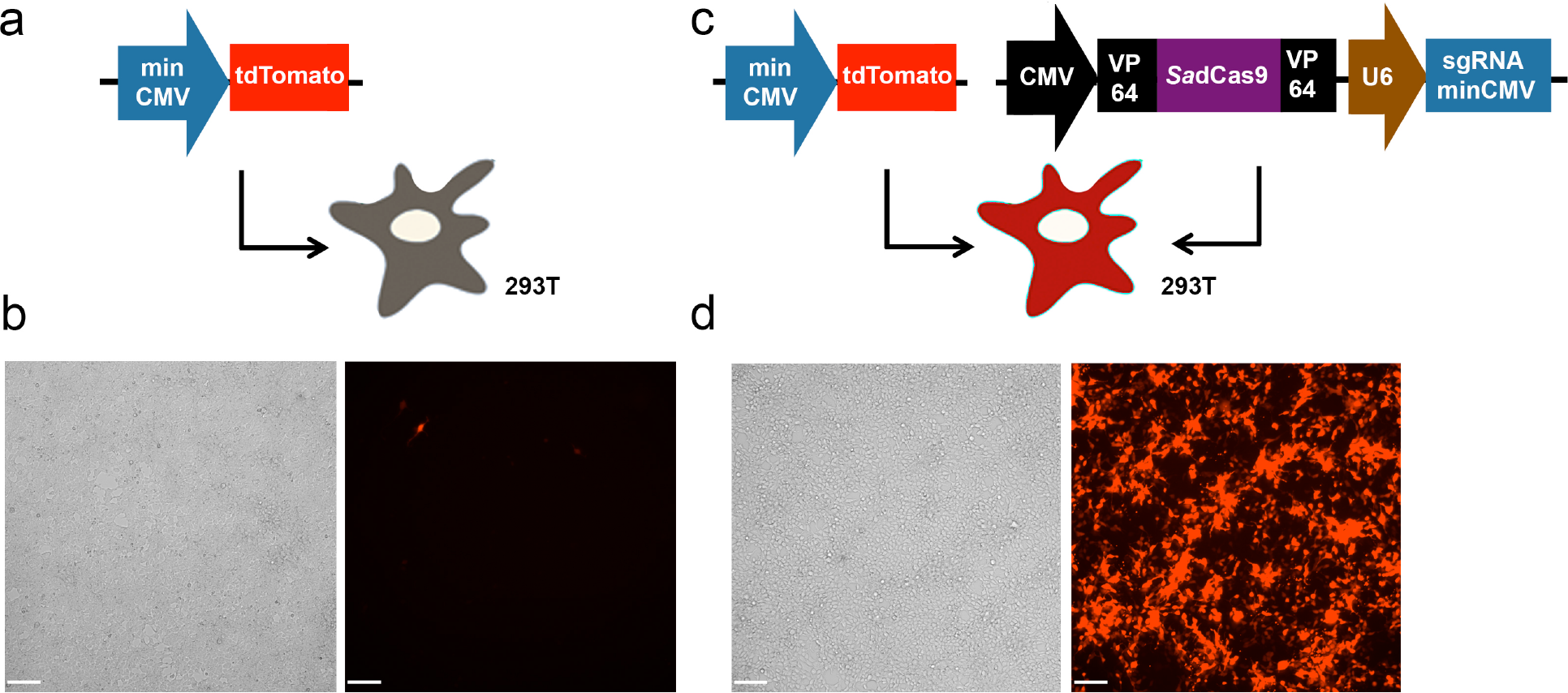
*Sa*dCas9-2xVP64 enhances expression of minCMV-driven *tdTomato in vitro*. HEK293T was transfected with (**a**,**b**) a plasmid containing minCMV-driven *tdTomato* gene only, or (**c**,**d**) in combination with a plasmid containing *Sa*dCas9-2xVP64 and a sgRNA targeting the minCMV promoter. (**b**,**d**) Cells were imaged for *tdTomato* expression by fluorescent microscopy. Scale bar: 50 μm.

**Fig. S2.**
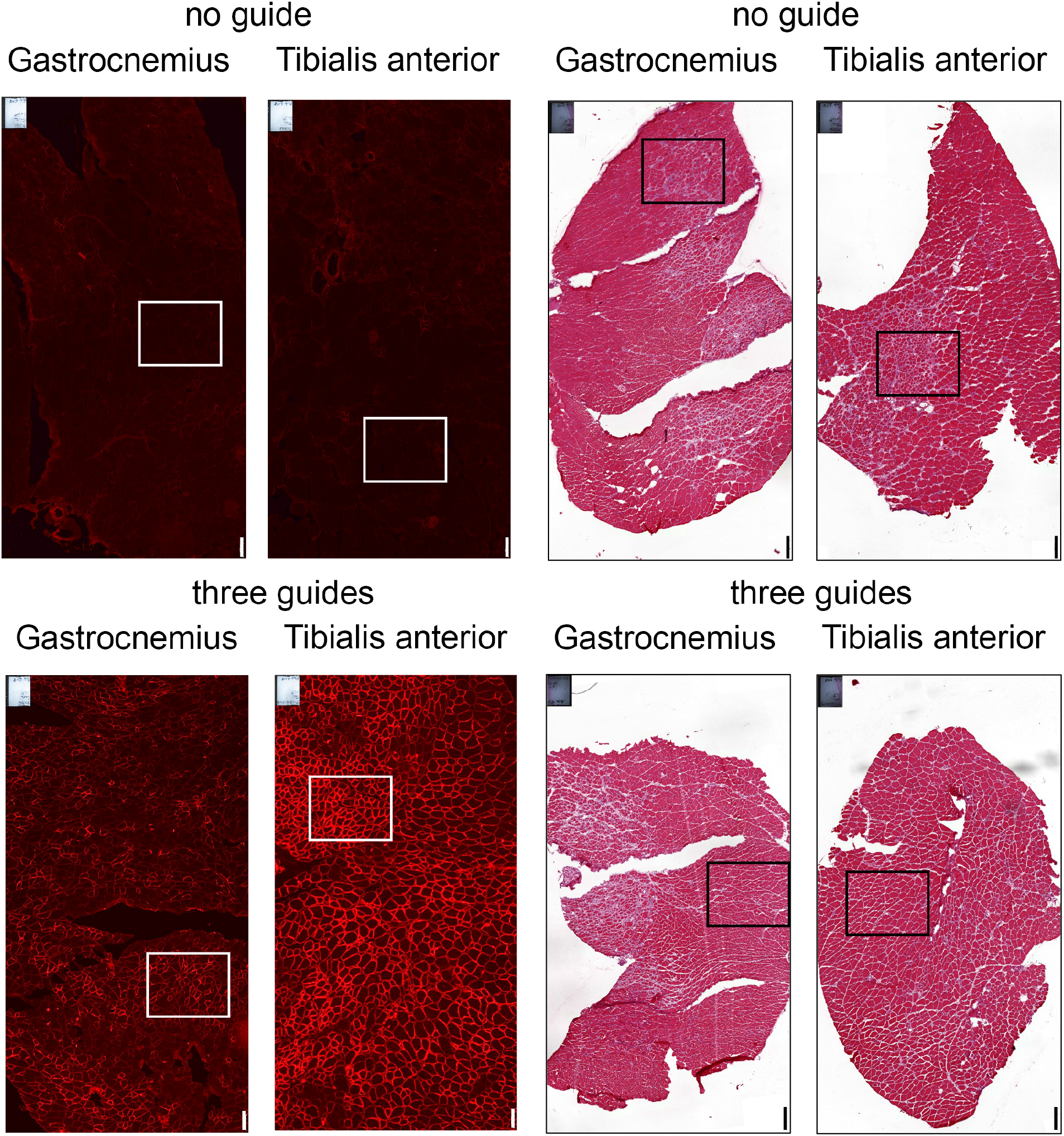
Expression of Lama1 and muscle architecture following early intervention in neonatal *dy^2j^/dy^2j^* mice. Lower magnification of immunofluorescence and H&E staining images of Fig. 3 in the main manuscript.

**Fig. S3.**
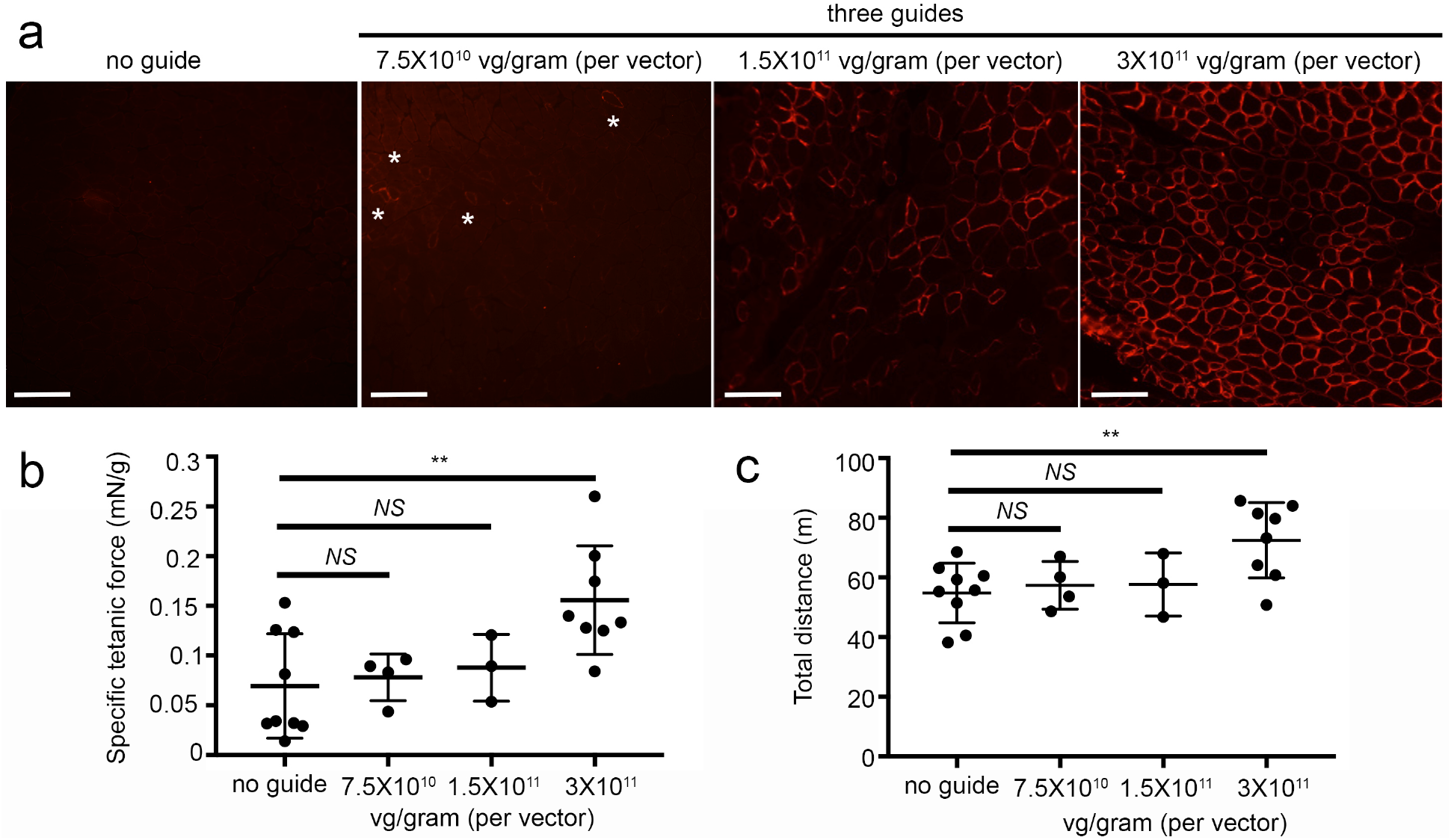
Upregulation of Lama1 corresponds to improvement of muscle functions. Three-weeks old *dy^2j^/dy^2j^* mice were injected systemically via tail vein with different doses of AAV9: 7.5×10^10^, 1.5×10^11^, or 3×10^11^ viral genome (vg)/gram of mouse. Two AAVs were necessary for the three guide cohort, therefore the total doses of virus injected was 1.5×10^11^, 3×10^11^, and 6×10^11^ viral genomes/gram of mouse. TA muscles isolated four weeks later were stained for Lama1 expression (**a**). Asterisks indicate Lama1-positive fibers in the low dose cohort. Scale bar: 100 μm. *In vivo* contractile force assay (**b**) and open field test (**c**) performed in the end of the treatment regimen. Data are presented as average ± standard deviation. Statistical analysis was performed using one-way ANOVA. *NS:* not significant. \*\**P<*0.01.

**Fig. S4.**
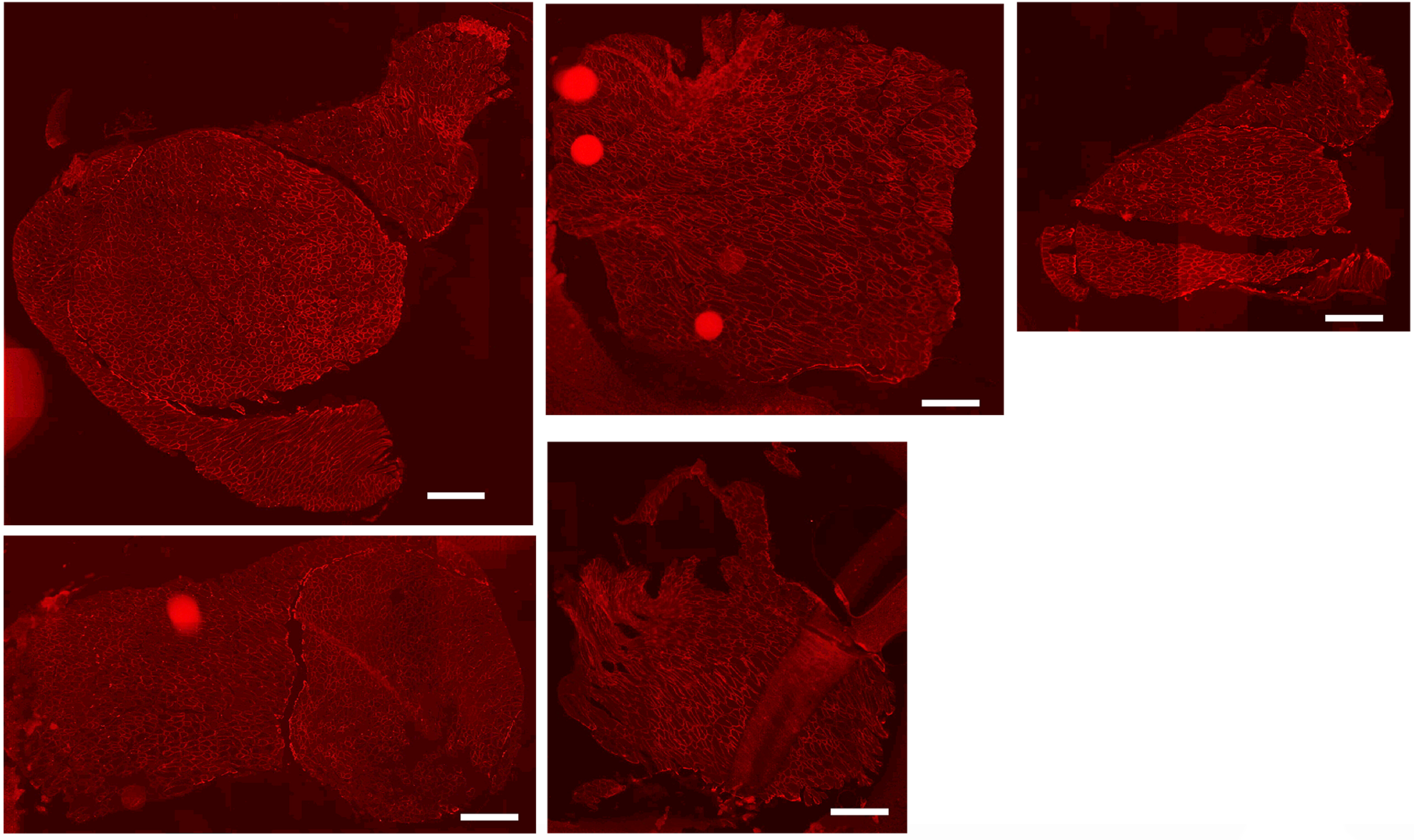
Representative images of Lama1-positive muscle sections. Mice were injected systemically with three guides at the age of three weeks old via tail vein, and sacrificed at the age of 11-12 weeks old. Muscles were stained for Lama1 expression (red). Scale bar: 500 μm.

**Fig. S5.**
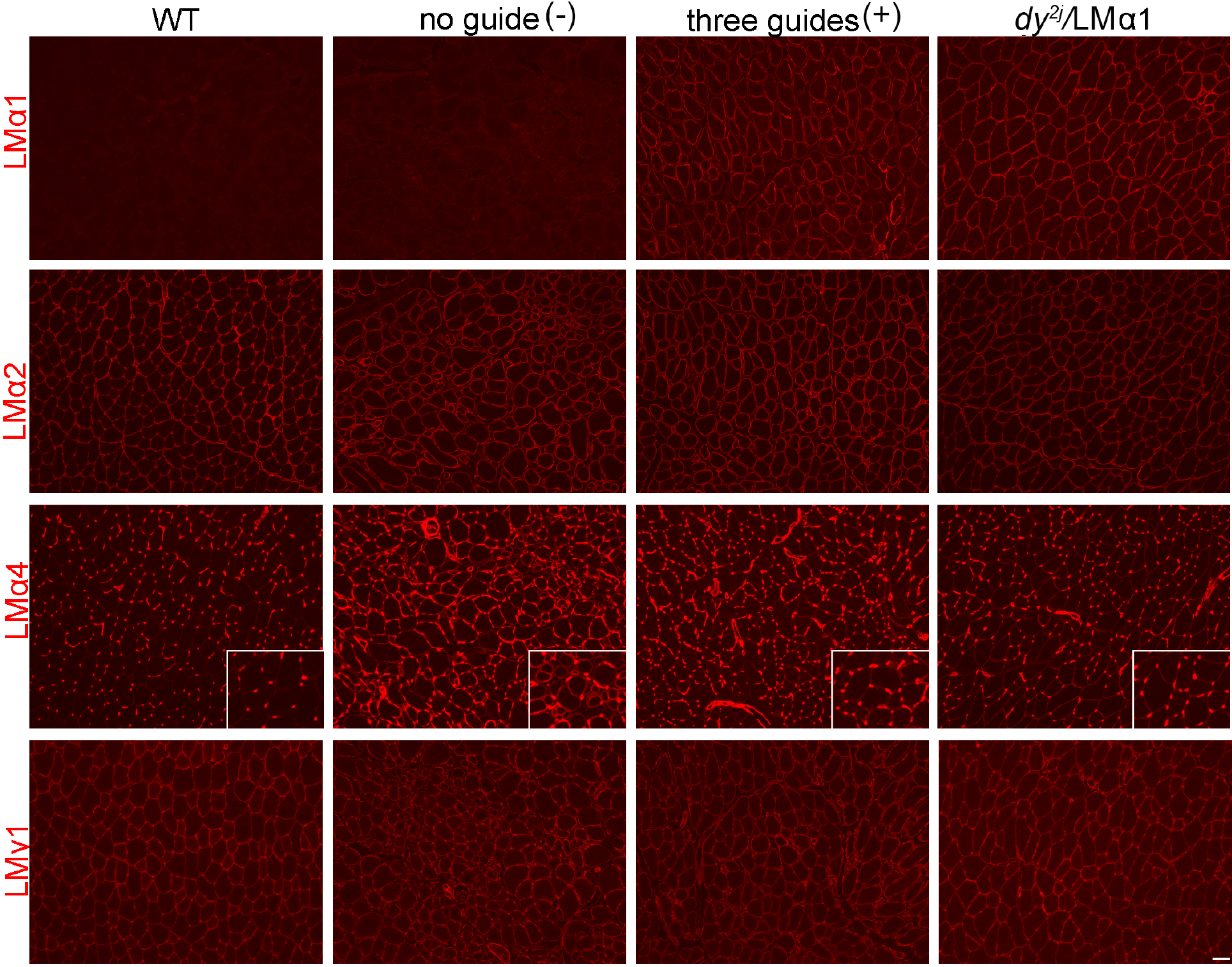
Expression of Laminin (LM) subunits in tibialis anterior muscles. Expression of LM subunits in tibialis anterior from wild-type, *dy^2j^/dy^2j^* (intravenously injected with AAV carrying no guide or three guides) and *dy^2j^/*LMα1 (with transgenic overexpression of LMα1) mice (n=3 for each group). Expression of LMα1 chain in tibialis anterior is comparable between *dy^2j^/dy^2j^* AAV-three guide-treated mice and transgenic mice. No major differences in expression of LMα2 and LMγ1 were detected between the groups. LMα4 chain is upregulated in muscle from *dy^2j^/dy^2j^* AAV-no guide-treated mice and its expression is partially normalized in muscle from *dy^2j^/dy^2j^* AAV-three guide treated mice. Scale bar: 50μm.

**Fig. S6.**
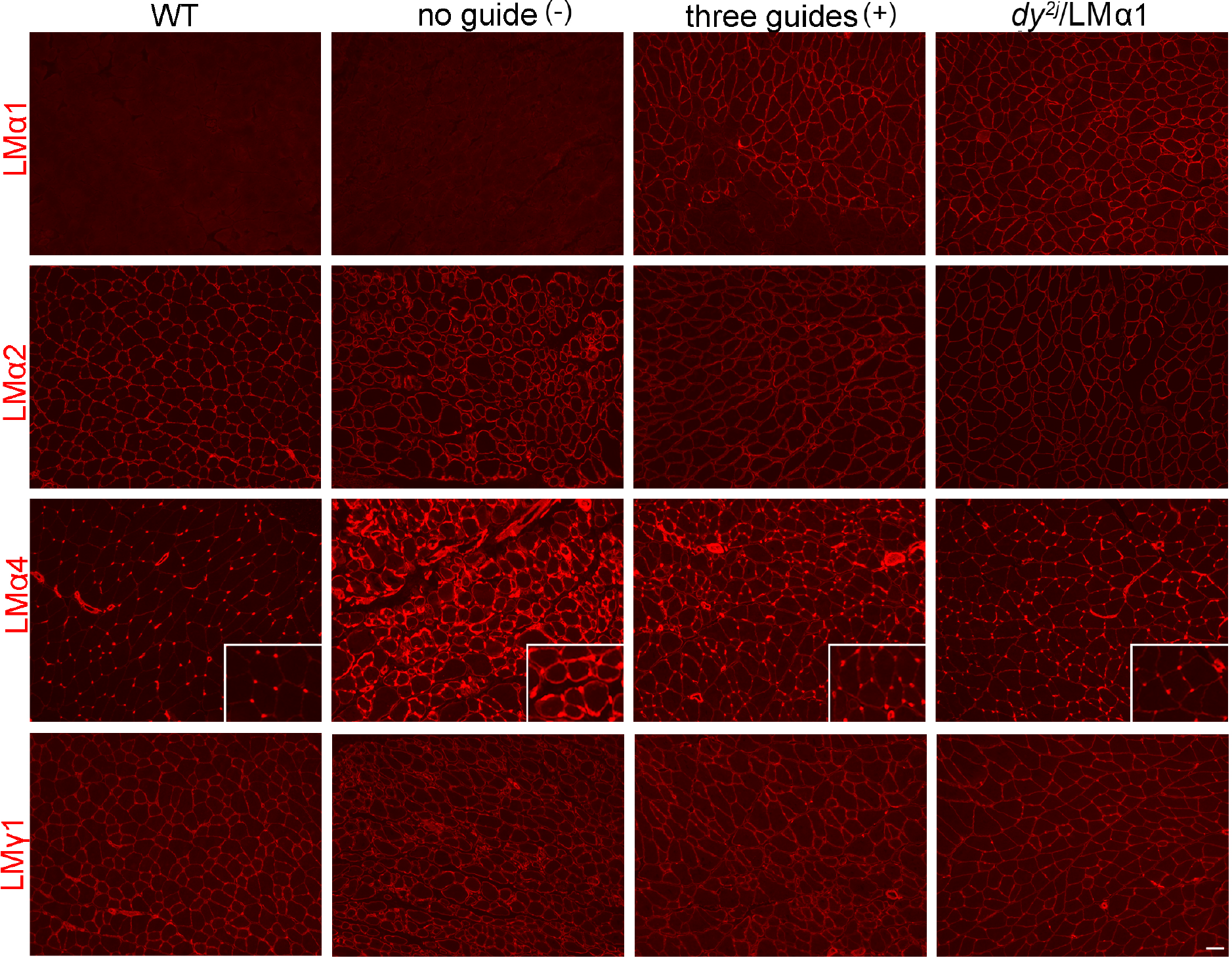
Expression of Laminin (LM) subunits in gastrocnemius muscles. Expression of LM subunits in tibialis anterior from wild-type, *dy^2j^/dy^2j^* (intravenously injected with AAV carrying no guide or three guides) and *dy^2j^/*LMα1 (with transgenic overexpression of LMα1) mice (n=3 for each group). Expression of LMα1 chain in tibialis anterior is comparable between *dy^2j^/dy^2j^* AAV-three guide-treated mice and transgenic mice. No major differences in expression of LMα2 and LMγ1 were detected between the groups. LMα4 chain is upregulated in muscle from *dy^2j^/dy^2j^* AAV-no guide-treated mice and its expression is partially normalized in muscle from *dy^2j^/dy^2j^* AAV-three guide treated mice. Scale bar: 50μm.

**Fig. S7.**
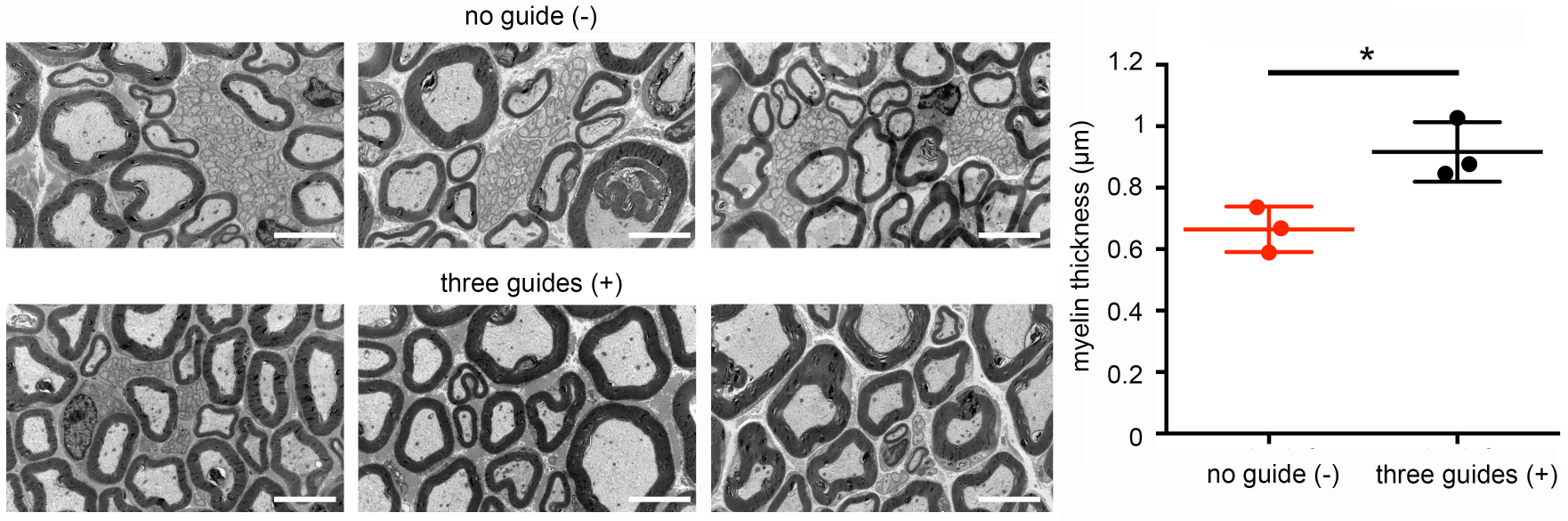
Myelination of sciatic nerves. Electron microscopy images of sciatic nerves isolated from *dy^2j^/dy^2j^* mice injected with AAV carrying no guide or three guides (n=3 for each group). Myelin thickness was quantified using ImageJ and presented as average ± standard deviation. Statistical analysis was performed using Student’s *t*-test. *\*P<*0.05. Scale bar: 5 μm.

**Fig. S8.**
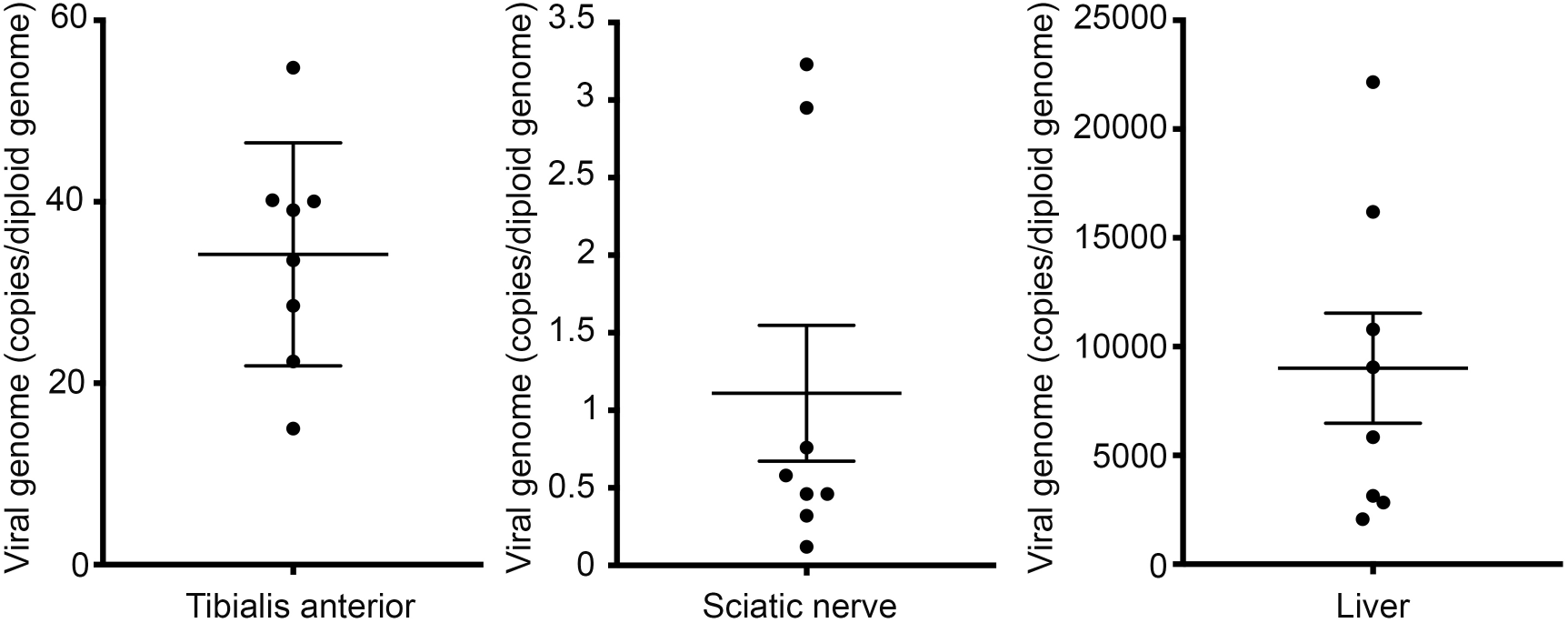
Quantitative evaluation of AAV genome distribution. Genomic DNA isolated from tibialis anterior muscle, sciatic nerve and liver of dy2j mice injected with AAV carrying three guides (n=8) was amplified for the presence of viral genome by qPCR. Data are presented as average ± SEM.

**Fig. S9.**
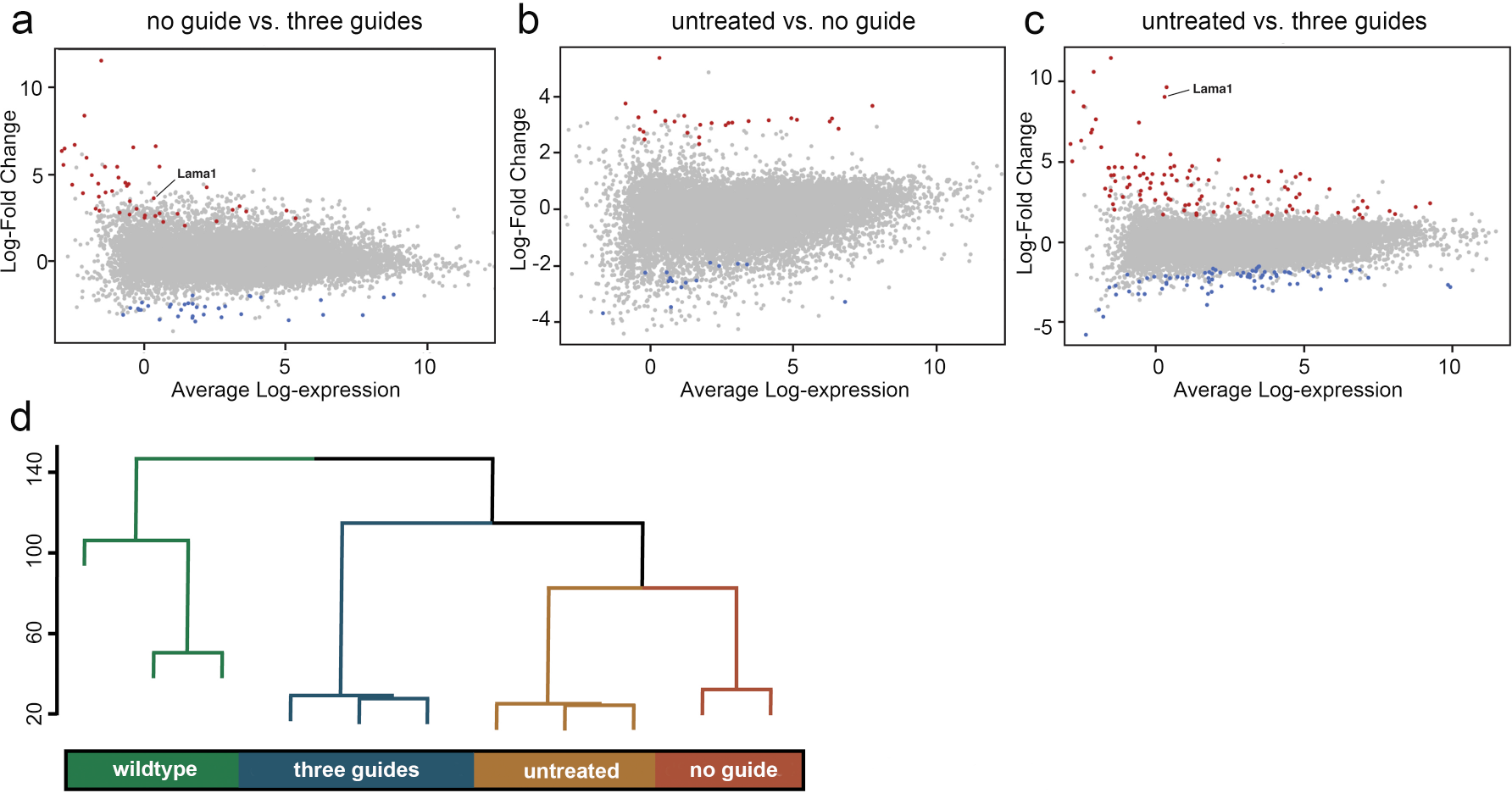
Genome-wide analysis of gene expression of *Sa*dCas9-VP64-treated *dy^2j^/dy^2j^* mice. Differential expression analysis derived from RNA-Sequencing results from the quadriceps of treated and untreated mice (**a-c**), comparing SadCas9-2xVP64 alone and SadCas9-2xVP64 with three guides targeting Lama1 (**a**), untreated to SadCas9-2xVP64 alone (**b**), and untreated to SadCas9-2xVP64 with three guides targeting Lama1 (**c**). Gene significantly differentially expressed, (FDR < 0.05), are coloured. Red data points indicate a log-fold change greater than one, while blue data points indicate a log-fold change less than one. Hierarchical clustering was performed on the normalized counts-per-million expression data (**d**).

**Fig. S10.**
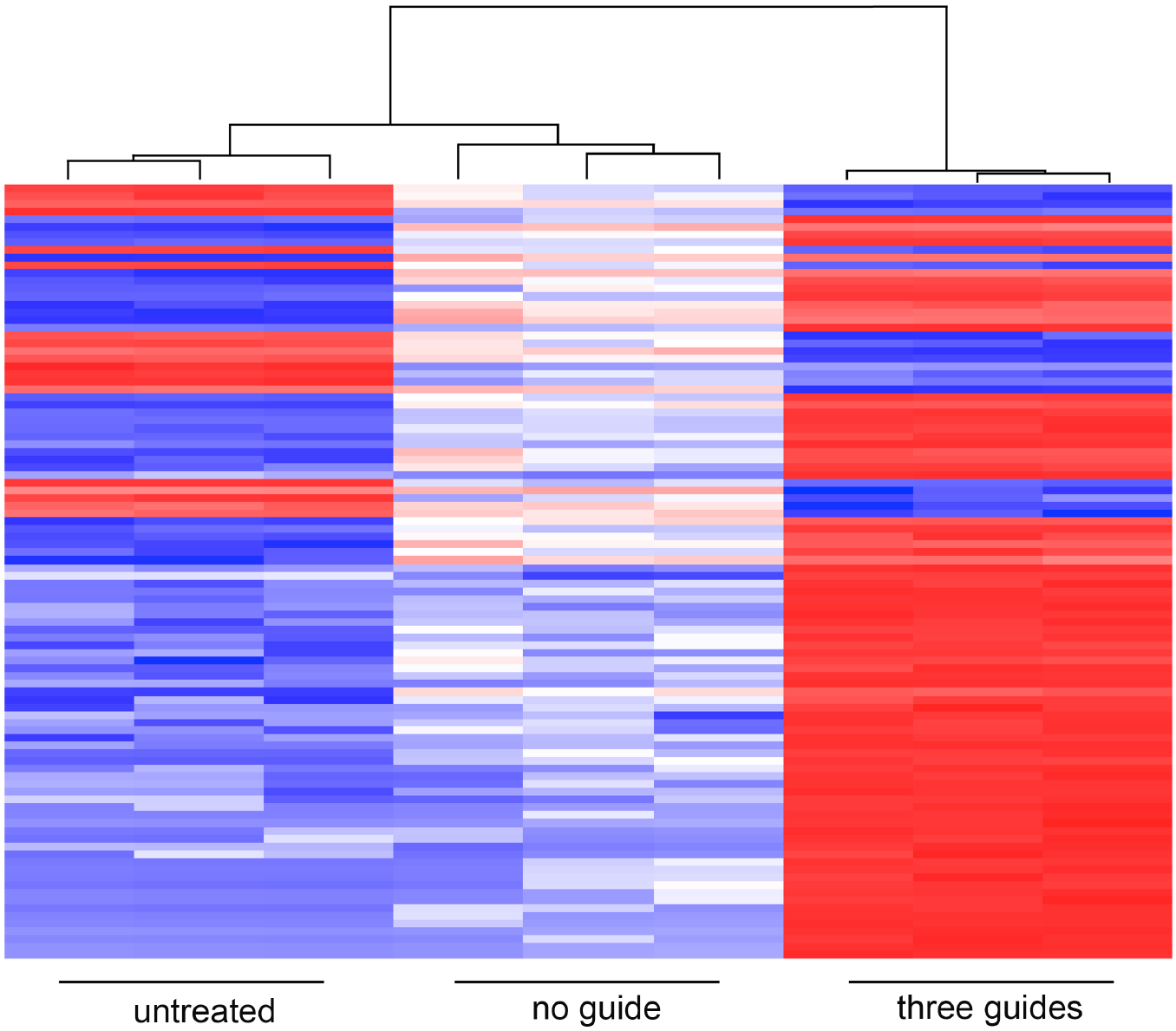
Top 100 differentially expressed genes. Heatmap illustrating normalized log-CPM values for the top 100 genes differentially expressed in SadCas9-2xVP64 with three guide treated quadriceps versus untreated quadriceps. Red indicates higher expression while blue indicates lower expression.

**Fig. S11.**
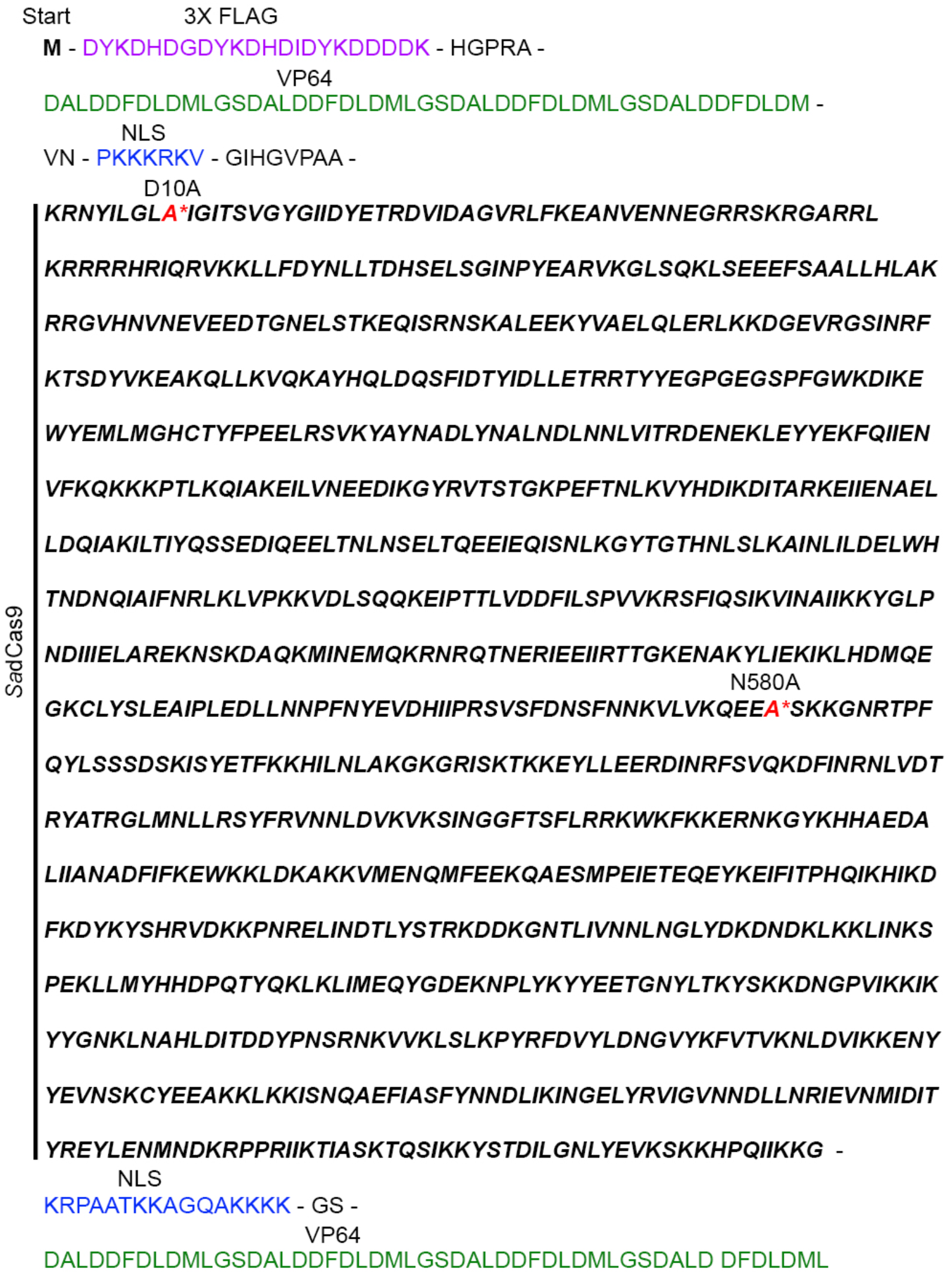
Protein encoded in the *Sa*dCas9-VP64 transcriptional activator plasmid. Different domains are annotated accordingly. Amino acids encoding *Sa*dCas9 are in bold and italic, and the mutated residues D10A and N580A are indicated in red asterisk (*). NLS: Nuclear localization signal.

## Supplemental video legends

**Video S1. Phenotype of control *dy^2j^/dy^2j^* mouse following early intervention.**

The *dy^2j^/dy^2j^* mouse was injected with AAV9 containing *Sa*dCas9-2xVP64 (no guide) at P2 (pre-symptomatic stage) via temporal vein and video was taken at the age of 7-week old. Hind limb paralysis, contracture and kyphosis resulting from lack of functional *Lama2* and compensatory *Lama1* are apparent.

**Video S2. Phenotype of treated *dy^2j^/dy^2j^* mouse following early intervention.**

The *dy^2j^/dy^2j^* mouse was injected with AAV9 containing *Sa*dCas9-2xVP64 and sgRNAs targeting *Lama1* proximal promoter (three guides) at P2 (pre-symptomatic stage) via temporal vein and video was taken at the age of 7-week old. Upregulation of compensatory *Lama1* expression ameliorates the hind limb paralysis, contracture and kyphosis.

**Video S3. Phenotype of control *dy^2j^/dy^2j^* mouse following intervention at symptomatic stage.**

The *dy^2j^/dy^2j^* mouse was injected with AAV9 containing *Sa*dCas9-2xVP64 (no guide) at 3-week old (pre-symptomatic stage) via tail vein. Video was taken at the age of 7-week old. Hind limb paralysis, contracture and kyphosis resulting from lack of functional *Lama2* and compensatory *Lama1* are apparent.

**Video S4. Phenotype of treated *dy^2j^/dy^2j^* mouse following intervention at symptomatic stage.**

The *dy^2j^/dy^2j^* mouse was injected with AAV9 containing *Sa*dCas9-2xVP64 and sgRNAs targeting *Lama1* proximal promoter (three guides) at 3-week old (pre-symptomatic stage) via tail vein. Video was taken at the age of 7-week old. Dystrophic features and disease progression were significantly improved and partially reversed following upregulation of *Lama1*.

## Supplemental tables

**Supplemental Table 1.**
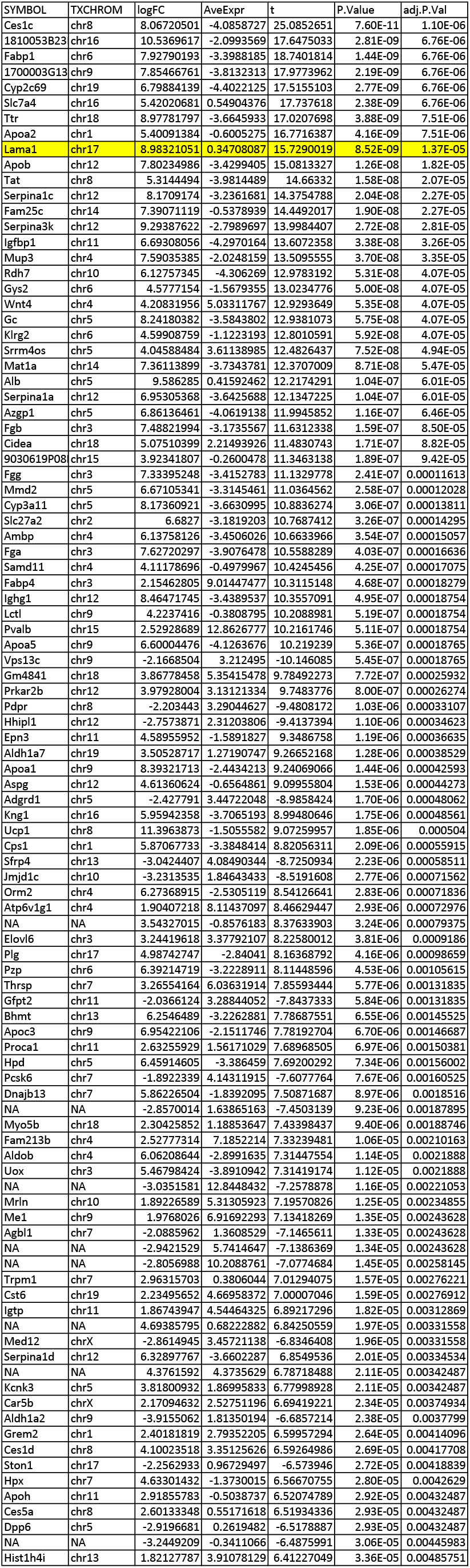
Top 100 genes differentially expressed in “SadCas9-2xVP64 + three guides” compared to “untreated”

**Supplemental Table 2.**
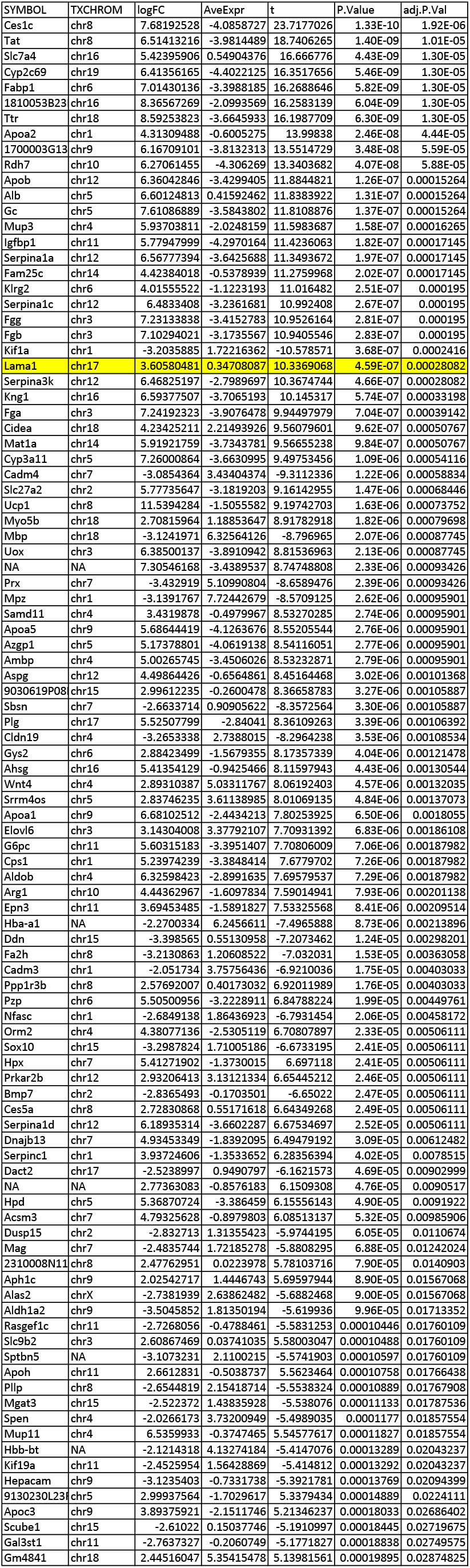
Top 100 genes differentially expressed in “SadCas9-2xVP64 + three guides” compared to “SadCas9-2xVP64 + no guide”

**Supplemental Table 3.**
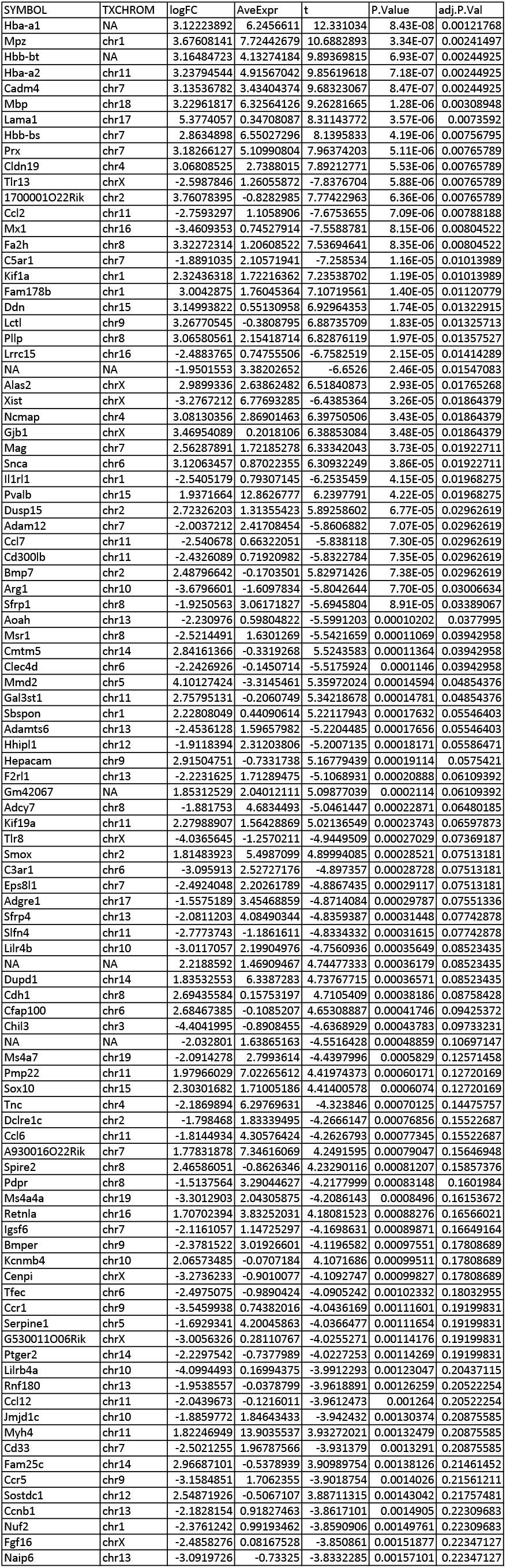
Top 100 genes differentially expressed in “SadCas9-2xVP64 + no guide” compared to “untreated”

**Supplemental Table 4.**
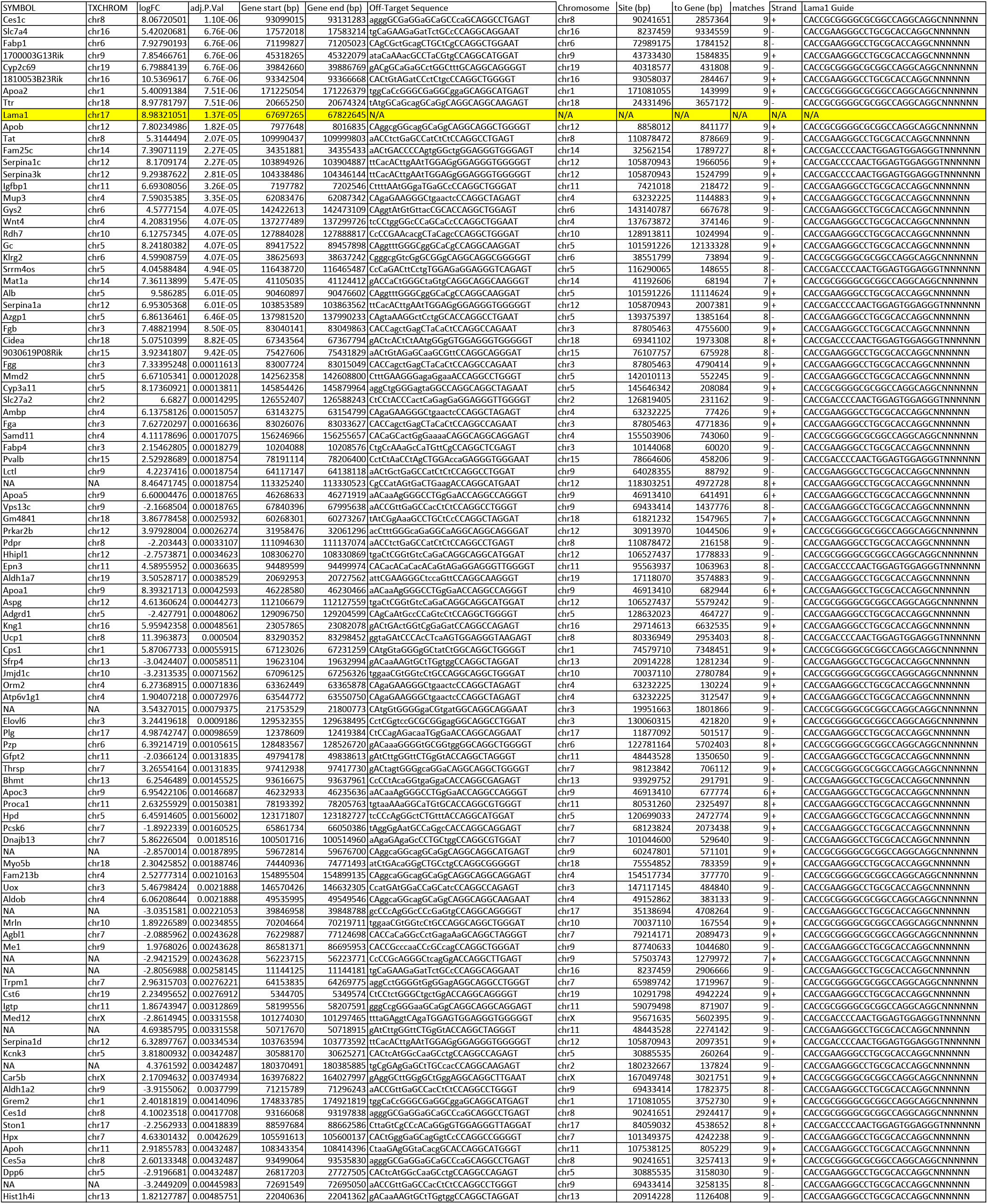
Analysis showing distance from off-target site to differentially expressed gene body

**Supplemental Table 5.**
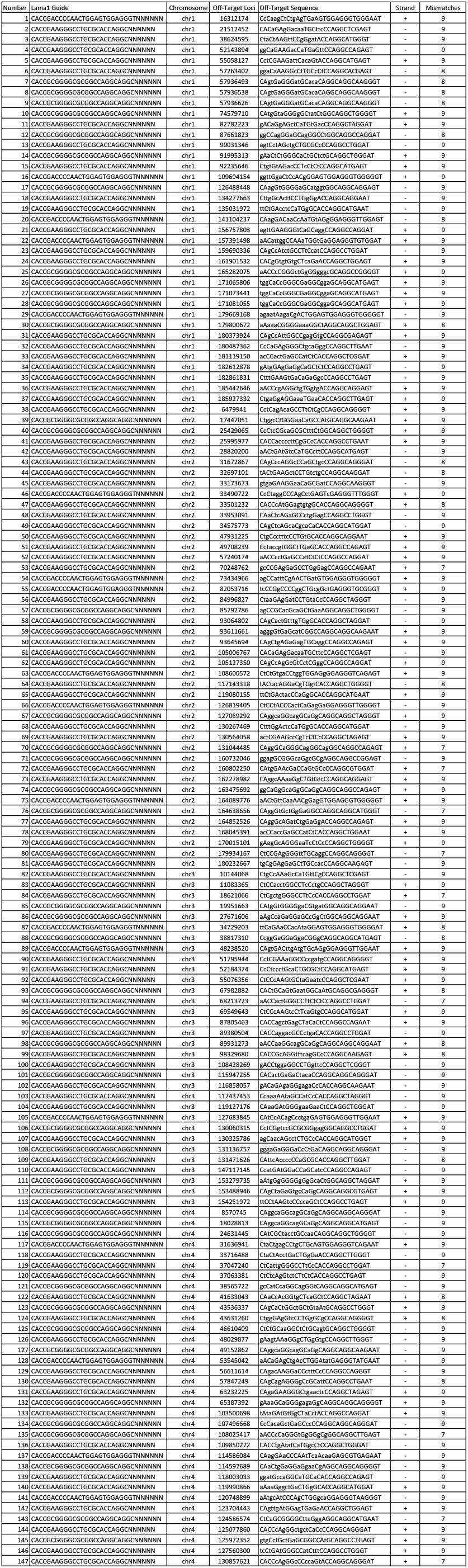
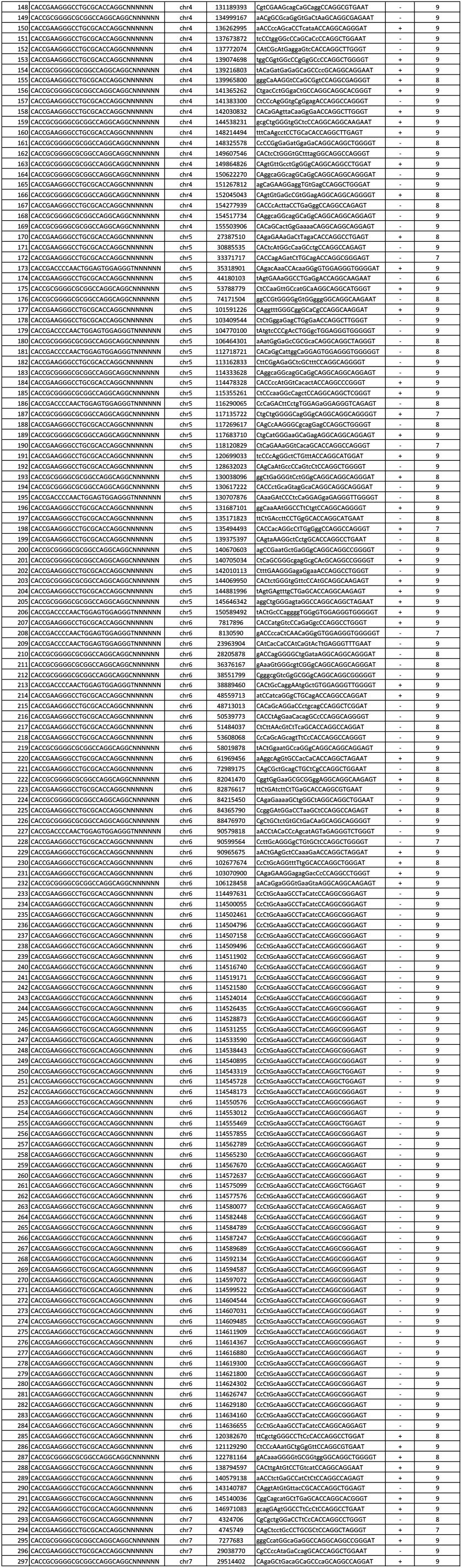
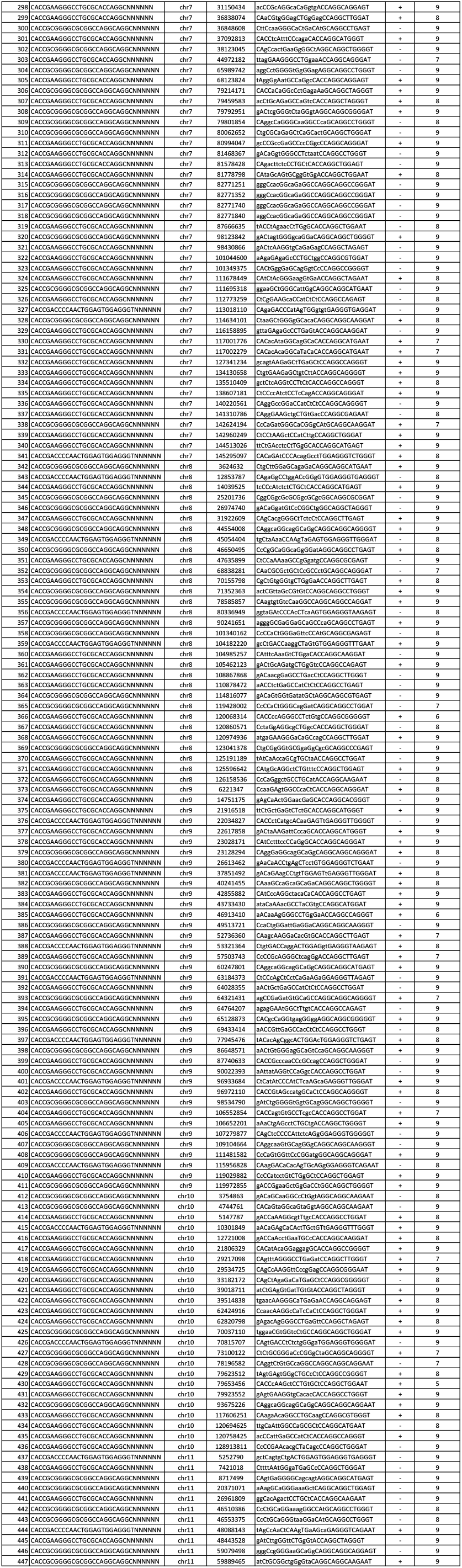
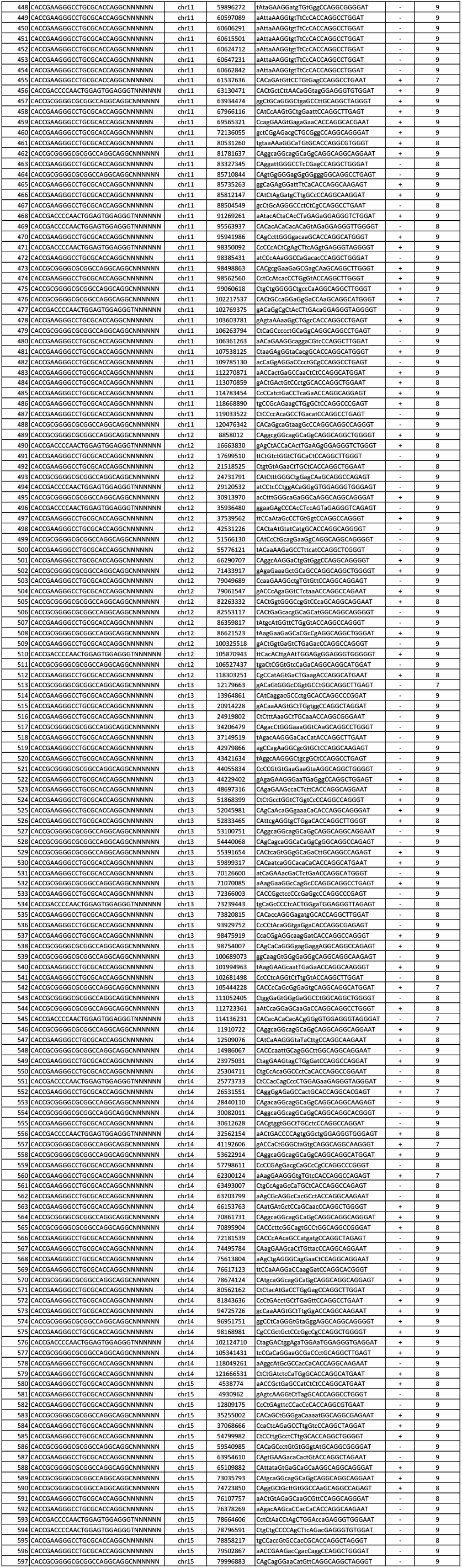
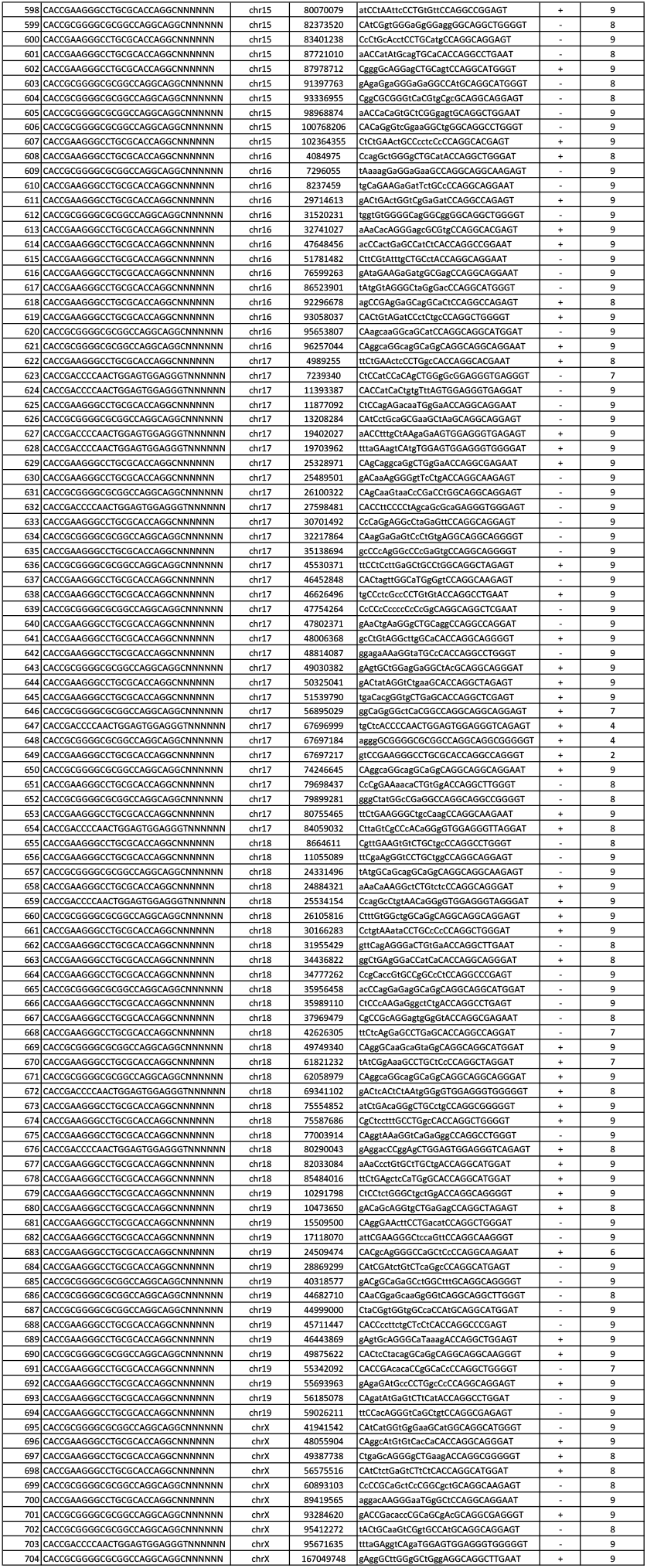
List of computationally predicted off-target sites

**Supplemental Table 6.**
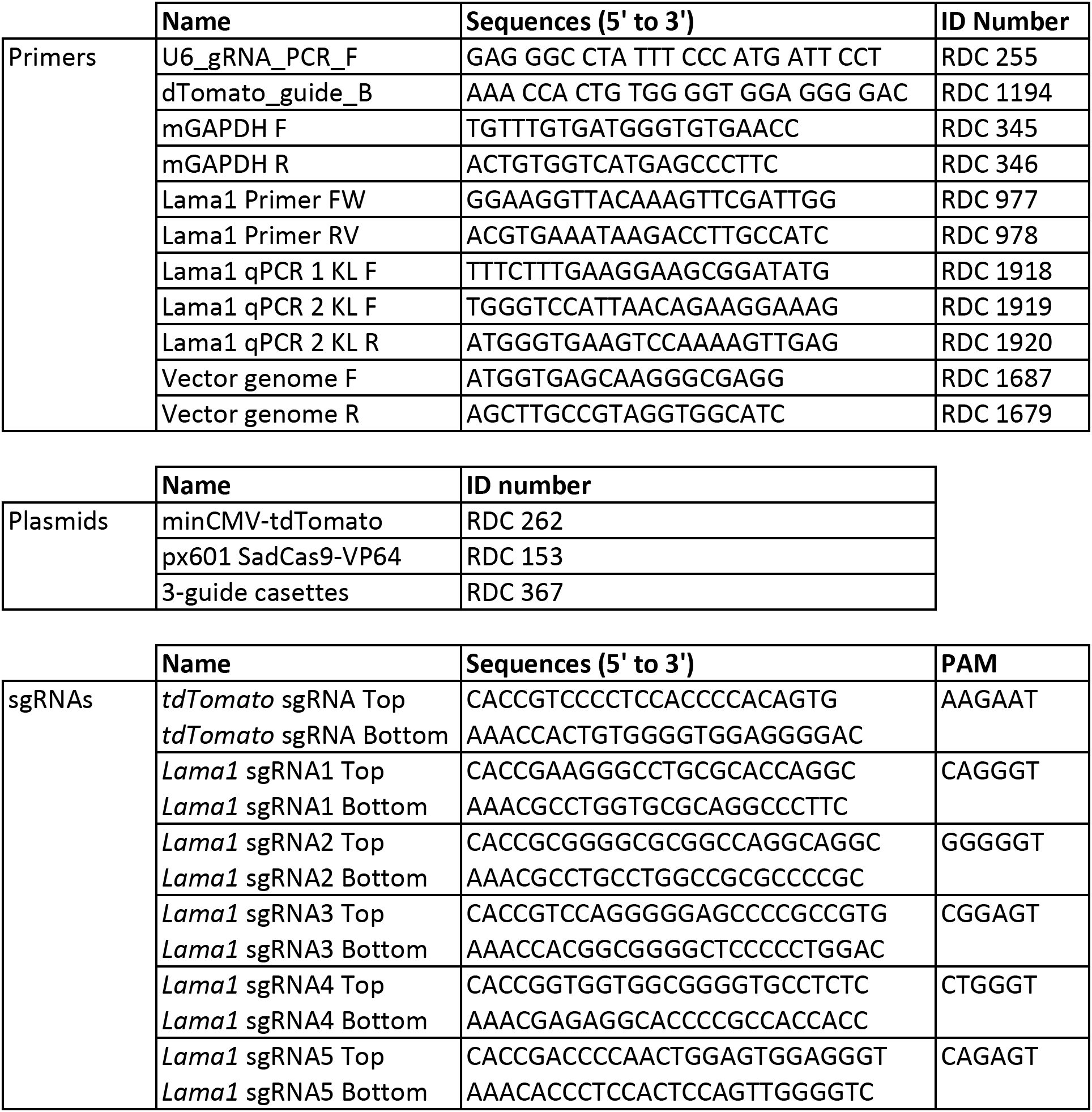
Primers, plasmids and sgRNAs used in this study

